# Identification and Characterization of Outer Membrane Proteins and Membrane Spanning Protein Complexes in *Brucella melitensis*

**DOI:** 10.1101/2025.07.27.666995

**Authors:** Jahnvi Kapoor, Amisha Panda, Ilmas Naqvi, Satish Ganta, Sanjiv Kumar, Anannya Bandyopadhyay

## Abstract

Brucellosis (Malta fever) is a zoonotic disease that affects both humans and animals, including cattle, sheep, and goats. *Brucella melitensis* is the most virulent and clinically significant species in humans. It is a Gram-negative bacterium with three groups of outer membrane proteins (OMPs): minor OMPs (Group 1), and major OMPs (Groups 2 and 3). OMPs with β-barrel architecture play important roles in nutrient transport, efflux, adhesion, and membrane biogenesis. Despite their importance, the structure, function, and interaction dynamics of several *B. melitensis* β-barrel OMPs and associated protein complexes remain mostly unexplored. In this study, we conducted a comprehensive in silico analysis to characterize known outer membrane β-barrel (OMBB) proteins and identify novel OMBBs in *B. melitensis* 16M. Proteins were modelled using five computational tools: AlphaFold 3, ESMFold, SWISS-MODEL, RoseTTAFold, and TrRosetta. Outer-membrane insertion of the novel OMBBs was confirmed using PPM 3.0, Protein GRAVY, DREAMM, and MemProtMD_Insane. Putative functions were predicted using structure- and sequence-based annotations. Sequence variation across 46 *B. melitensis* strains were identified and mapped onto the structural models. OMBB-associated protein complexes – the RND (Resistance-Nodulation-Division) efflux pumps, the lipopolysaccharide transport (Lpt) complex, and the β-barrel assembly machinery (BAM) complex – were modelled, and protein–protein interactions (PPIs) were analyzed to confirm thermodynamically stable assemblies. This study presents a robust in silico strategy for exploring OMP architecture and provides valuable structural insights to support the development of diagnostics, targeted therapeutics, and vaccines against *B. melitensis*.

## 1. Introduction

Brucellosis (Malta fever) is a zoonotic infection caused by various *Brucella* spp. that afflicts both livestock and humans. Worldwide, the annual burden is estimated at 1.6–2.1 million new human cases,^1^ with the disease persisting in resource-limited regions in Africa, Central and South America, Asia, Eastern Europe, the Mediterranean Basin, and the Middle East.^2,3^ Clinical manifestations range from intermittent fever, fatigue, and arthralgia to severe sequelae such as arthritis, hepatitis, spontaneous abortion or premature birth.^2,4^ Transmission occurs through direct contact with infected animal tissues or fluids (e.g., aborted material, vaginal discharge), ingestion of unpasteurized dairy products, inhalation of contaminated aerosols in abattoirs and meat-processing plants, and, less frequently, intrauterine transmission from an infected mother to her unborn child during pregnancy.^1,2,5,6^ *Brucella* are small (0.5–0.7 μm × 0.6–1.5 μm), aerobic, Gram-negative, non-motile coccobacilli^7^ with an incubation period ranging from 5 days to 6 months.^8^ *Brucella melitensis* is the most virulent species. Upon ingestion, *B. melitensis* passes through the stomach and crosses epithelial barriers to disseminate systemically.^4^ It is carried by macrophages to lymphoid tissues, spreading through the lymphatic system.^2^ Because its symptoms are non-specific, brucellosis is frequently misdiagnosed.^9^ Serological assays such as Buffered Acidified Plate Antigen Test (BAPAT), Rose Bengal Plate Test (RBPT), and Milk Ring Test (MRT) remain the mainstays of diagnosis,^10–12^ while doxycycline and rifampin are commonly used antibiotics for treatment. Despite the veterinary use of live-attenuated vaccines (e.g., *B. abortus* S19, RB51, *B. melitensis* Rev1), no licensed human vaccine exists.^13^

*B. melitensis* biovar 1 strain 16M is the principal agent of ovine and caprine brucellosis and a significant human pathogen.^2^ *B. melitensis* was the first *Brucella* species whose genome was sequenced, and subsequent proteomic maps have been produced for strains 16M and Rev1.^14–16^ *B. melitensis* 16M serves as the NCBI reference with two circular chromosomes totaling 3.295 Mb and encoding 2981 proteins.^17^ Like other Gram-negative bacteria, *B. melitensis* possesses an inner phospholipid membrane, a periplasmic peptidoglycan layer, and an outer membrane (OM) whose inner leaflet contains phospholipids and whose outer leaflet is rich in lipopolysaccharide (LPS).^18^ *Brucella* spp. encodes multiple families of outer membrane proteins (OMPs), including many heat-stable porins and lipoproteins. The bacterium encodes numerous OMPs, classified into three groups: the minor OMPs of Group 1 (88-94 kDa) and major OMPs of Group 2 (porin, 36–38 kDa) and Group 3 (25–27 kDa and 31-34 kDa).^19^ Minor OMPs of Group 1 consist of lipoproteins Omp10 and Omp19. Major OMPs of Group 2 comprise two closely related porins, Omp2a and Omp2b. These are 16 stranded β-barrel proteins with large surface-exposed loops. Group 3 consists of two families of eight-stranded β-barrel proteins - Omp25 and Omp31. Group 3 proteins are crucial virulence determinants that mediate host-cell interactions, for example, Omp25 of *Brucella abortus* binds the dendritic-cell receptor SLAMF1, dampening inflammatory cytokine release.^20,21^ *B. melitensis* LPS is structurally atypical compared with enterobacterial LPS; its smooth phenotype hinders phagosome maturation and oxidative bursts.^22^ LPS is delivered to OM by the Lpt complex, yet the key OM component LptD remains poorly characterized in this species- an important gap given its role in immune evasion. In addition, *B. melitensis* secretes outer-membrane vesicles (OMVs) laden with OMPs, LPS, lipoproteins, and periplasmic factors that potently stimulate innate immunity; 29 OMV proteins (52% OM-derived) were catalogued in strain 16M.^23,24^ Consequently, elucidating the structure, function, and biogenesis of OMPs is essential for diagnostics, vaccine design, and therapeutics.

In the present study, we applied a consensus-based computational pipeline to identify outer-membrane β-barrel (OMBB) proteins in *B. melitensis* 16M. Twenty-four OMBBs were identified and grouped according to the extent of their prior characterization (Table 1). High-confidence structural models for each protein were generated with five independent modelling tools, and putative functions were assigned through complementary structure- and sequence-based annotations. Mapping sequence polymorphisms from 46 *B. melitensis* strains onto the models revealed genetic diversity, evolutionary pressures, and variable surface regions. Among the OMBB proteins were homologues of TolC, LptD, and BamA- signature components of the RND efflux pumps, Lpt (lipopolysaccharide-transport) complex, and β-barrel assembly machinery (BAM) complex, respectively. We reconstructed the complete Bam, Lpt, and two tripartite RND efflux systems (BepDE-BepC and BepFG-BepC) and delineated protein-protein interfaces by identifying inter-subunit hydrogen bonds. Structure-guided alignments with *E. coli* BAM complex allowed us to predict interaction hotspots in BamA that are critical for OM assembly and thus potential targets for antimicrobial intervention.

**Table 1:** Grouping of identified OMBB proteins based on earlier studies.

| **Groups of OMBB proteins** | **Criteria for Grouping** | **Accession No. (Protein name<sup>†</sup>)** |
| --- | --- | --- |
| **Group A** | Proteins having experimental OMP evidence and in silico structural models | WP_002964666.1 (OmpW family protein)<br>WP_002965367.1 (Omp25)<br>WP_002965368.1 (Omp25)<br>WP_004683466.1 (Omp25)<br>WP_004683739.1 (Omp25)<br>WP_004681766.1 (Omp31)<br>WP_002970988.1 (Omp2a)<br>WP_006256196.1 (Omp2b) |
| **Group B** | Proteins having experimental OMP evidence but lack structural predictions | WP_002966799.1 (Multidrug efflux RND transporter BepC)<br>WP_004681095.1 (TonB-dependent receptor domain-containing protein)<br>WP_004683979.1 (TonB-dependent receptor domain-containing protein)<br>WP_005972352.1 (TonB-dependent receptor domain-containing protein) |
| **Group C** | Proteins having only in silico studies | WP_002971090.1 (hypothetical protein)<br>WP_004683433.1 (LPS-assembly protein LptD)<br>WP_004683866.1 (Outer membrane protein assembly factor BamA)<br>WP_004683944.1 (Outer membrane protein)<br>WP_004684144.1 (Outer membrane protein)<br>WP_004684330.1 (YadA-like family protein) |
| **Group D** | Novel OMBB proteins, merely annotated in NCBI or UniProt without experimental or computational characterization | WP_002968232.1 (outer membrane beta-barrel protein)<br>WP_004682315.1 (outer membrane protein)<br>WP_004684108.1 (alginate export family protein)<br>WP_004684499.1 (autotransporter outer membrane beta-barrel)<br>WP_005970295.1 (autotransporter assembly complex family protein)<br>WP_075589504.1 (OmpP1/FadL family transporter) |

<sup>†</sup>Protein names from NCBI, searched using Protein Accession no.

**Table 2:**
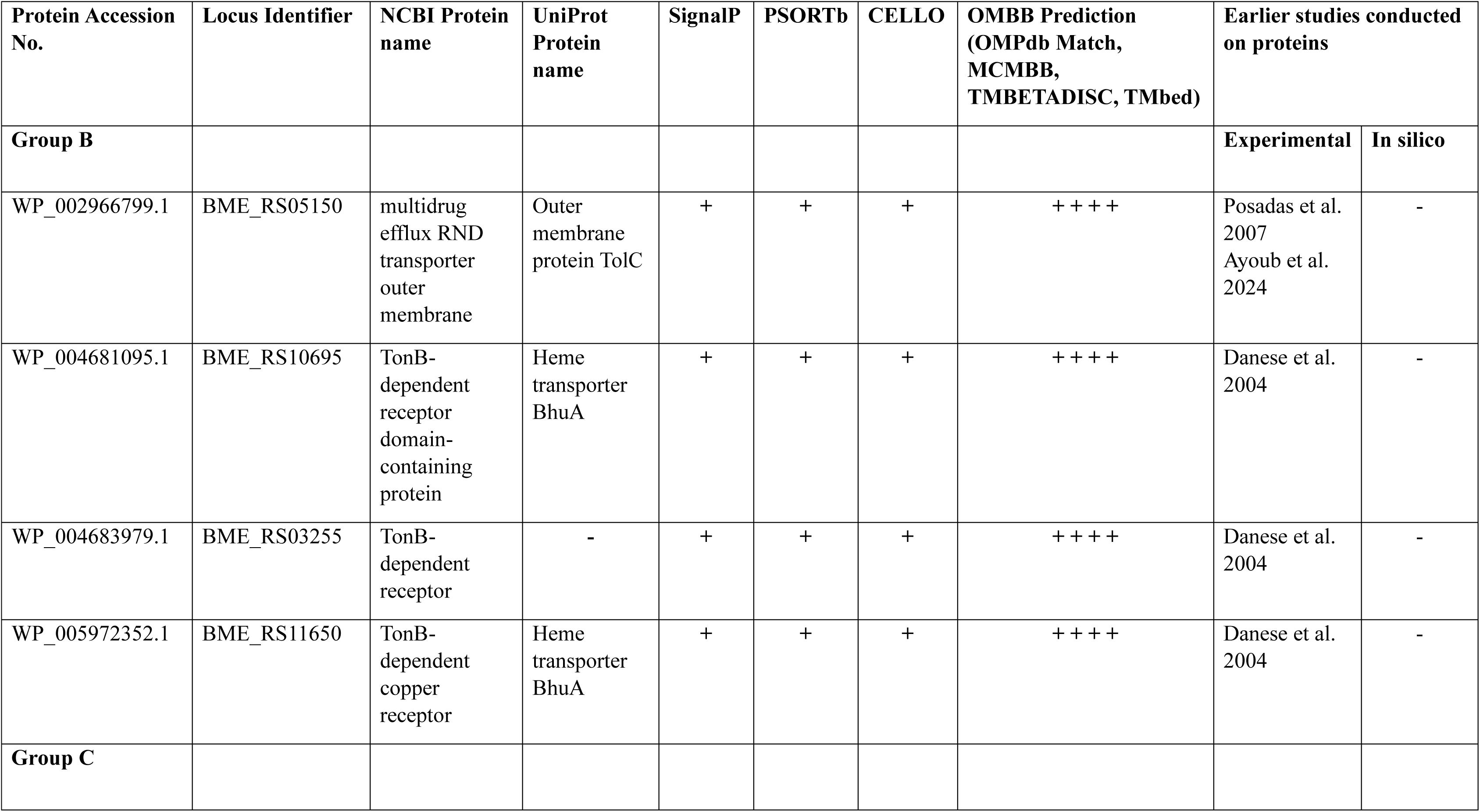
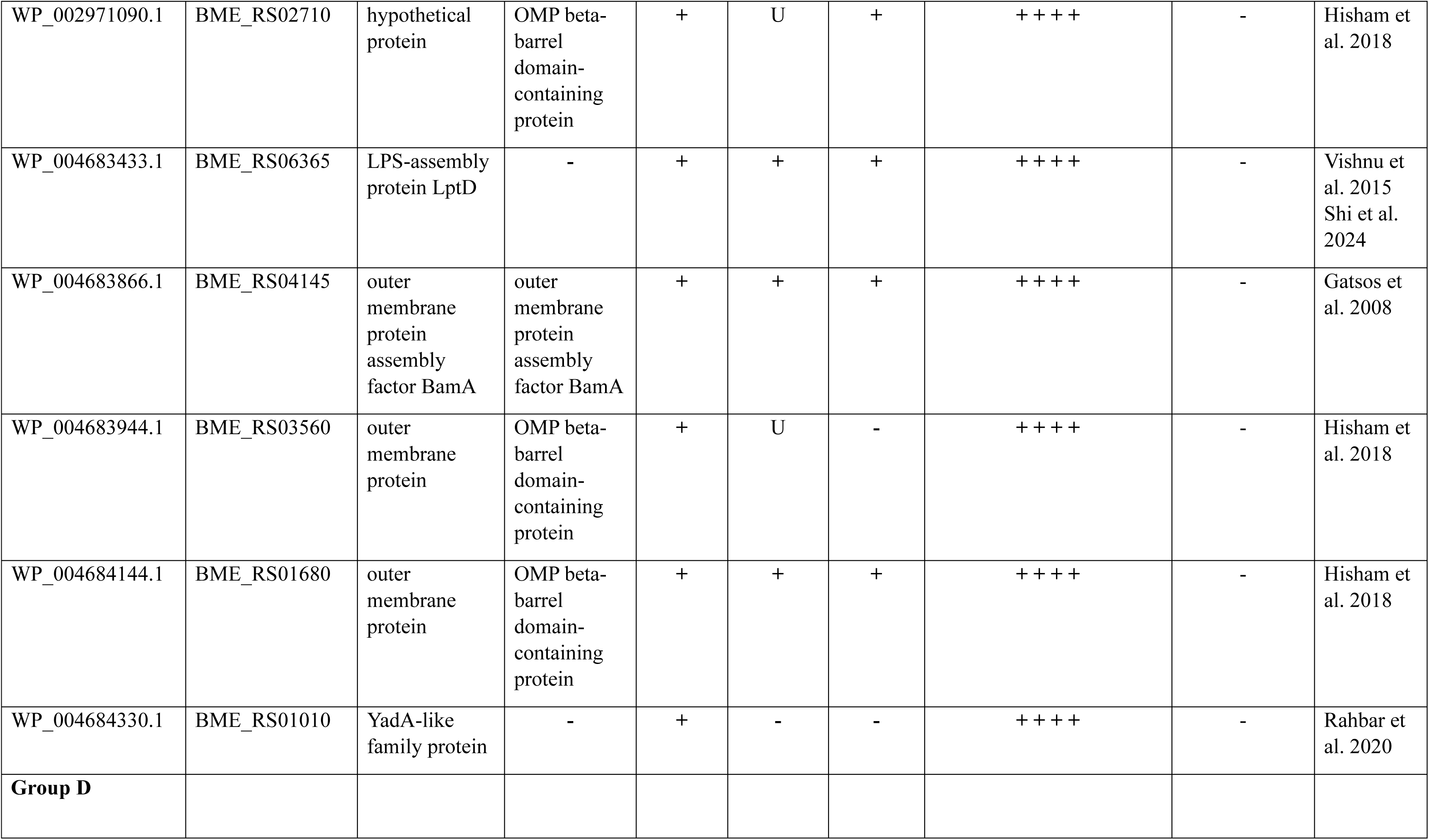
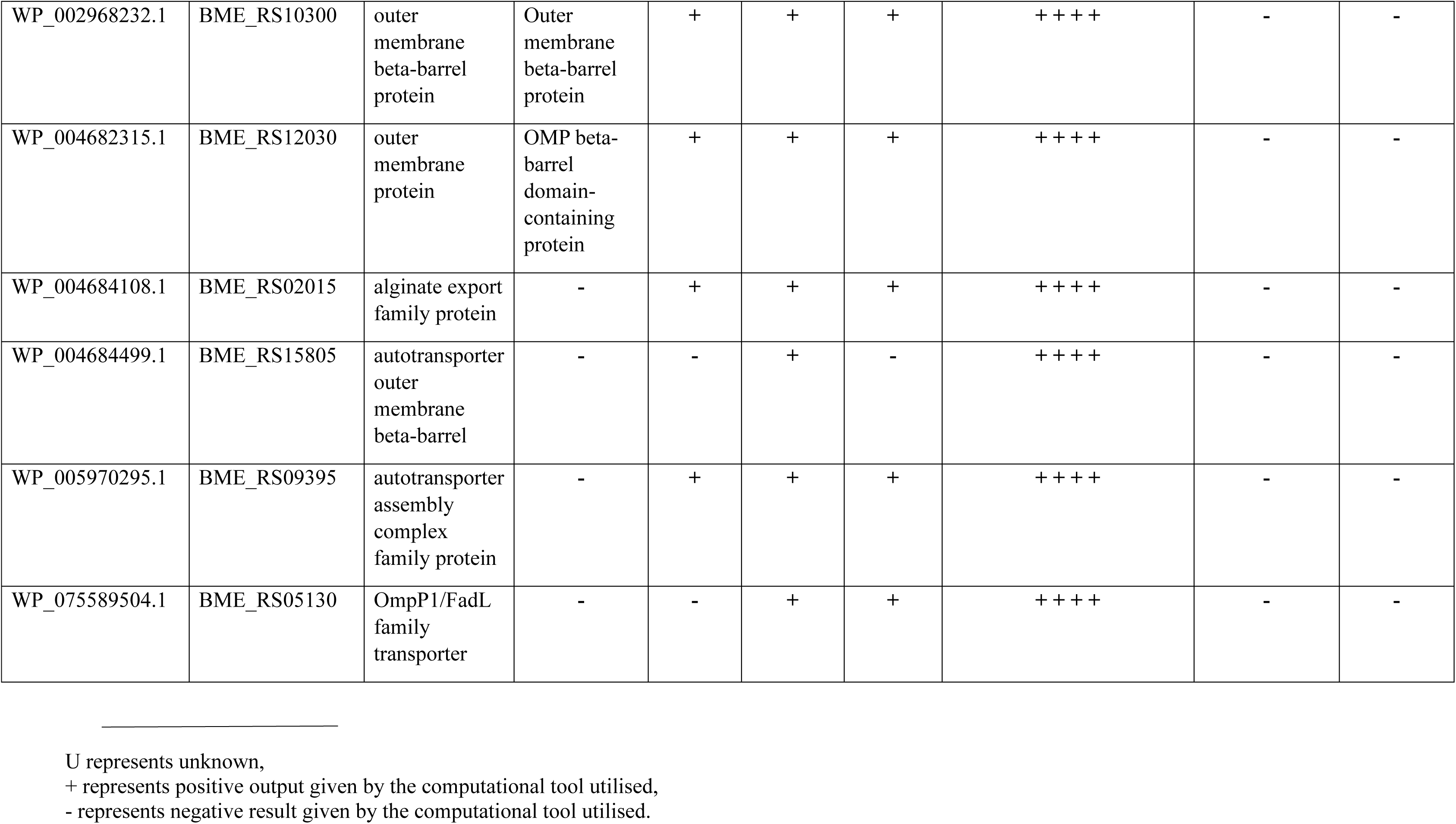
Predicted outer membrane β-barrel proteins from *B. melitensis* 16M.

## 2. Materials and Methods

### 2.1 Prediction of outper membrane β-barrel proteins

*B. melitensis* bv1 strain 16M, a reference strain, was selected for our study. Amino acid sequences of all the proteins of *B. melitensis* 16M were downloaded from NCBI^25^ using genome assembly ASM74041v1 (Accessed on 21 September 2024).

Nine bioinformatic tools were used to select OMPs. Subcellular localization was predicted using CELLO v.2.5^26^ and PSORTb 3.0^27^, providing insights into their cellular distribution. SignalP 5.0 was used to detect signal peptides and predict their cleavage sites.^28^ Physicochemical properties- peptide length, molecular weight, net charge, and isoelectric point (pI)- were determined with Pepstats tool from the EMBOSS package.^29^ SPAAN was used to predict adhesins and adhesin-like proteins.^30^ Functional domains within the protein sequences were identified with the Conserved Domain Database (CDD) (all tools accessed on 23 September 2024).^31^

The OMP selection pipeline is detailed below and schematically represented in Figure 1. We used a consensus-based computational approach to identify OMBB proteins. Outputs from four OMBB prediction tools were considered- one database OMPdb^32^, and three prediction tools - MCMBB^33^, TMBETADISC-RBF^34^, and TMbed^35^ (all tools accessed on 27 September 2024). Proteins that were predicted to have a β-barrel architecture by all four tools were considered for further investigation (Table S1). Additional 48 proteins were also identified as potential OMBB proteins by three of the four OMBB prediction tools but they were excluded due to lower confidence in their predictions (Table S6).

**Figure 1:**
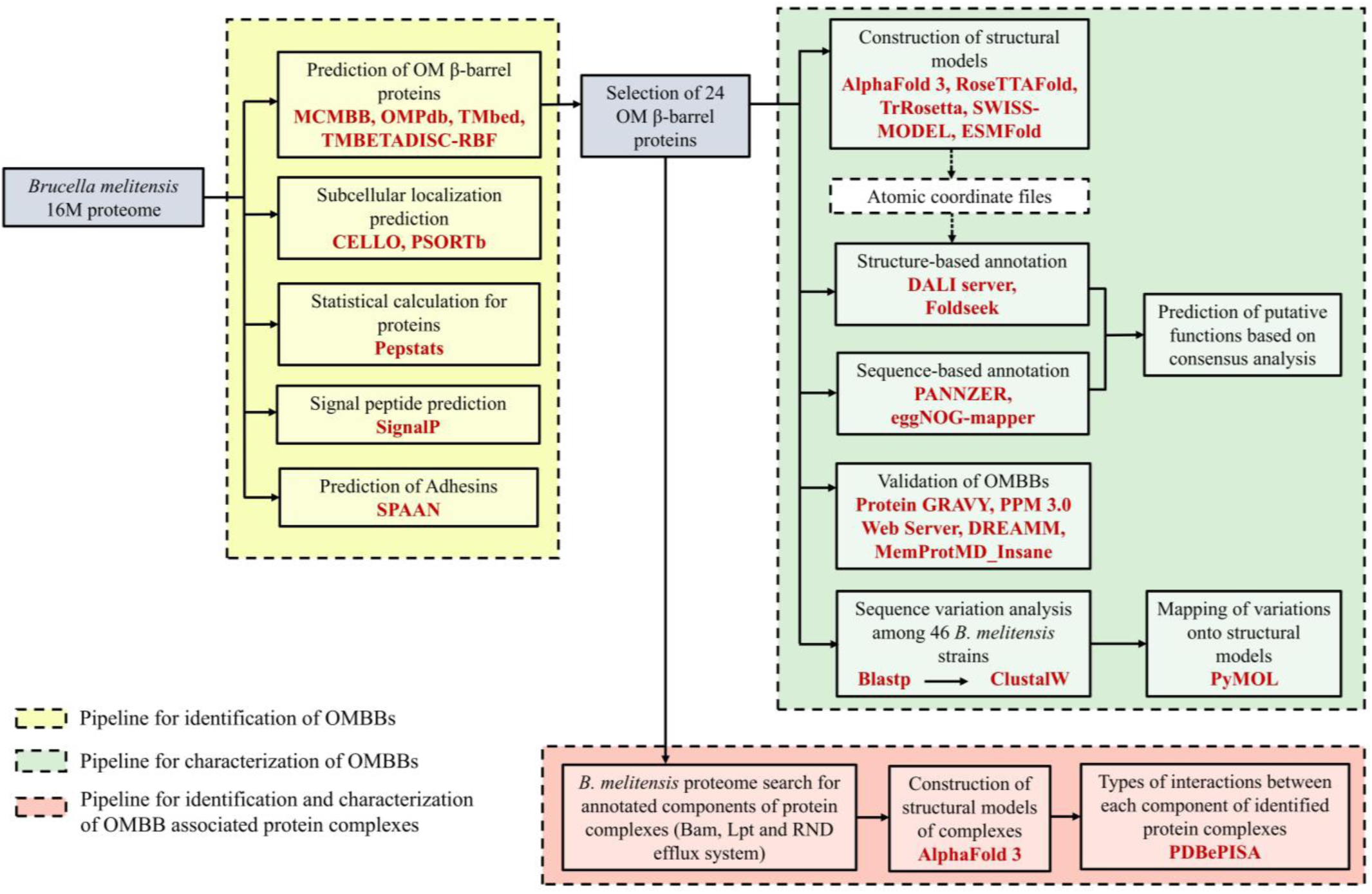
Computational framework for prediction, validation and characterization of OM β-barrel proteins from *B. melitensis* 16M. *B. melitensis* proteome was analyzed using tools such as CELLO, PSORTb, OMPdb, MCMBB, Tmbed, TMBETADISC-RBF, SignalP. Size, charge, and isoelectric point were predicted using Pepstats and SPAAN was employed to predict adhesin-like proteins. Structural models were generated using five tools: AlphaFold 3, RosettaFold, TrRosetta, SwissModel and ESMfold. Coordinate files of these structural models generated from AlphaFold 3 were used as queries in the DALI server and Foldseek, and putative functions were annotated. Sequence-based annotation was performed using PANNZER and eggNOG-mapper. Membrane insertion of identified proteins was validated using Protein GRAVY, PPM 3.0 web server, DREAMM and MemProtMD_Insane. Amino acid sequence variation analysis across 46 *B. melitensis* strains was performed using ClustalW and mapped onto structural models using PyMOL. A proteome-wide search identified annotated components of OM biogenesis and efflux complexes, followed by structure modelling using AlphaFold 3. Also, the interfacing residues and the type of interactions between different components of each protein complex was predicted.

### 2.2 Structural modelling of OMBB proteins

Three-dimensional structures of predicted proteins were generated from AlphaFold 3 (accessed on 10 October 2024).^36^ AlphaFold includes confidence metrics: pLDDT (predicted local distance difference test), PAE (predicted aligned error), pTM (predicted template modelling), and ipTM (inter-chain predicted TM) scores. Although the pLDDT scores were high, we modelled each protein with four additional tools- ESMFold^37^, SWISS-MODEL^38^, RoseTTAFold^39^, and TrRosetta^40^ (all tools accessed on 15 October 2024)- to enhance confidence in the structural predictions. For proteins that did not form complete barrels, we explored the possibility of oligomeric assemblies, prioritizing models with high pLDDT scores. Top-ranked models were selected on the basis of highest confidence scores and used for further analysis. Atomic coordinates were visualized in PyMOL 3.0.^41^ Structural similarity among the five models of each protein was assessed with US-align^42^ online web server and the corresponding RMSD (Root Mean Square Deviation) values were used to validate the predicted β- barrel architecture (Table 3). A low RMSD value reflects greater confidence in the accuracy and reliability of the predicted structures. Because all aligned structures had RMSD < 10 Å, we proceeded with the AlphaFold 3 models for further study.

**Table 3:**
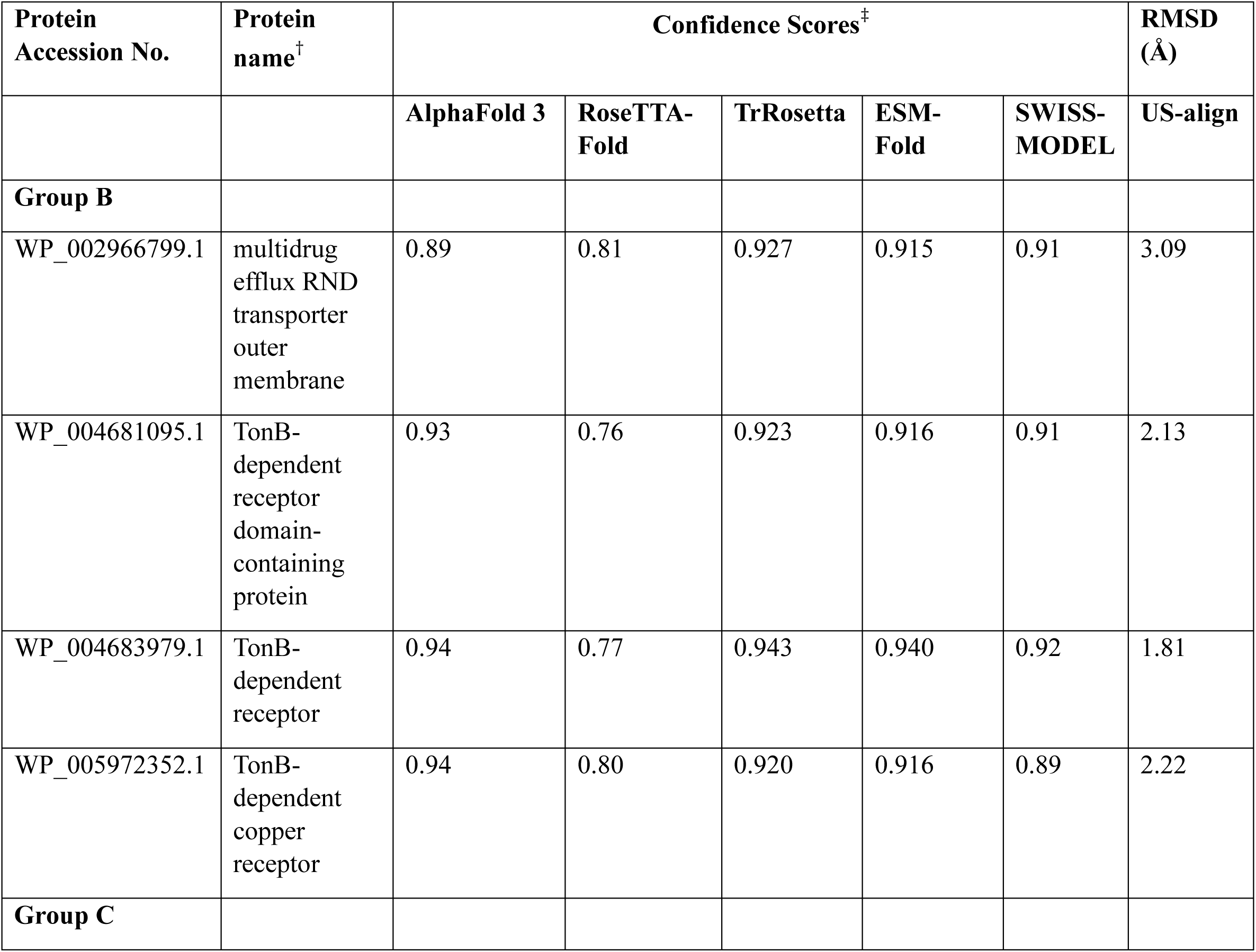
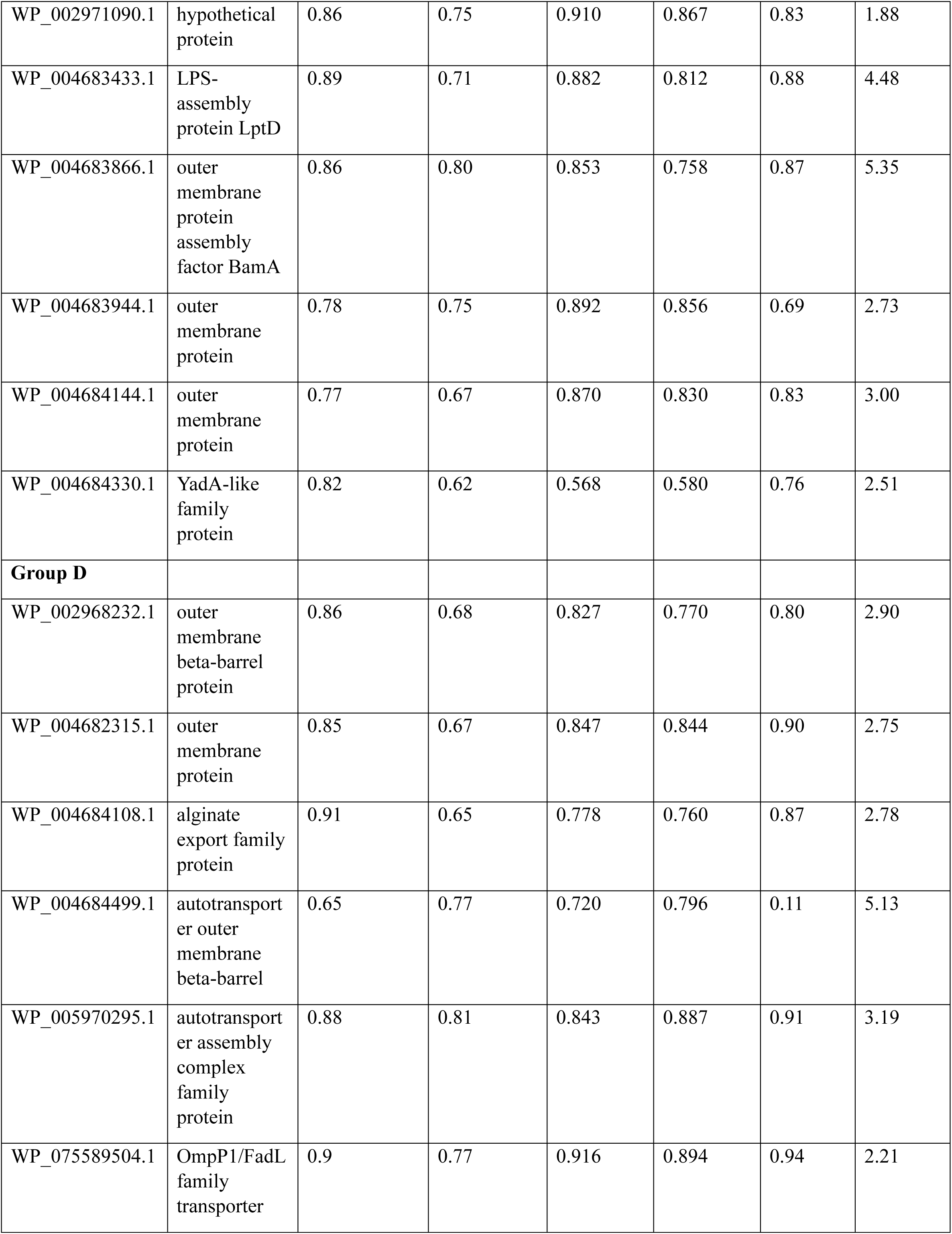
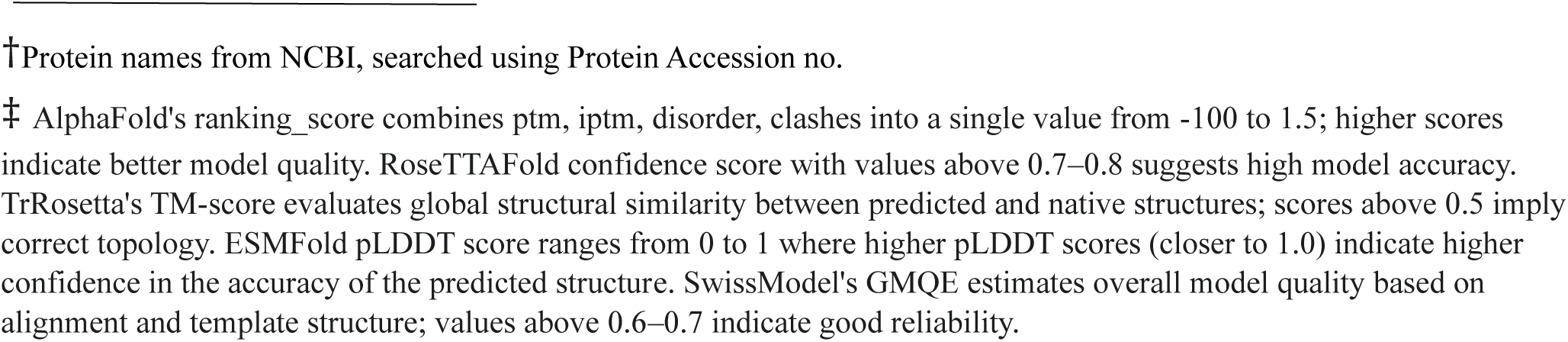
Structural alignment of 3D models of the identified proteins generated by five modelling tools.

### 2.3 Validation of outer membrane insertion

We further evaluated the uncharacterized novel OMBB proteins using four tools- Protein GRAVY^43^, DREAMM^44^, PPM 3.0 web server^45^ and MemProtMD_Insane^46^ (all tools accessed on 20 March 2025)- to assess their potential insertion into the outer membrane. Protein GRAVY calculates the GRAVY (grand average of hydropathy) score for the protein sequences by adding the hydropathy value for each residue and dividing by sequence length. The GRAVY value was used to predict hydrophobic regions of proteins which are likely to interact with outer membrane. DREAMM (Drugging pRotein mEmbrAne Machine learning Method) predicts protein-membrane interfaces using ensemble machine learning. PPM 3.0 determines optimal spatial positions of proteins in membranes and calculates the ΔG_transfer energy, the free energy required to transfer a protein from an aqueous environment into the lipid bilayer. MemProtMD_Insane streamlines the process of embedding membrane proteins into lipid bilayers for MD simulations. It uses INSANE (INSert membrANE) to construct realistic lipid bilayer environments tailored to the protein’s requirements, automating the placement of lipids around the protein to mimic its native membrane environment. We simulated each protein in a bacterial-like outer membrane (outer leaflet: 7 POPE, 2 POPG; inner leaflet: 7 POPE, 2 POPG, 1 cardiolipin) for a 500 ns production run. Results were visualized in PyMOL, and backbone RMSD calculated after alignment to the initial structure. Low RMSD values indicate high protein stability in the membrane.

### 2.4 Structure- and sequence-based functional annotation

Since some proteins were unannotated in NCBI or UniProt, we used structure-based approach to infer putative functions of the predicted β-barrel proteins. Atomic coordinates of the models were submitted to the DALI server^47^ (full PDB search option) to compare against all structures in the Protein Data Bank (PDB). The protein with the highest Z-score was taken as the top functional hit, and its reported function was retrieved from the literature. For a comprehensive and robust analysis, we also used Foldseek^48^ to search five major structure databases (AFDB-Proteome, AFDB-SWISSPROT, AFDB50, CATH50, and PDB100). The match with the highest TM-score was selected to assign the most likely function.

Sequence-based annotation complemented these analyses. PANNZER (Protein ANNotation with Z-scoRE)^49^, a tool designed to annotate proteins of prokaryotic or eukaryotic origin based on sequence data, was used for functional annotation. PANNZER evaluates the accuracy of predicted Gene Ontology (GO) classes using Positive Predictive Value (PPV), which measures the reliability of the annotations. The protein sequences were also submitted to eggNOG-mapper^50^, which utilizes the eggNOG database to assign orthologous groups, GO terms, KEGG pathways, COG functional categories, and predicted protein domains. This combined pipeline enabled consensus functional annotation of the query sequences (Table 4).

**Table 4:**
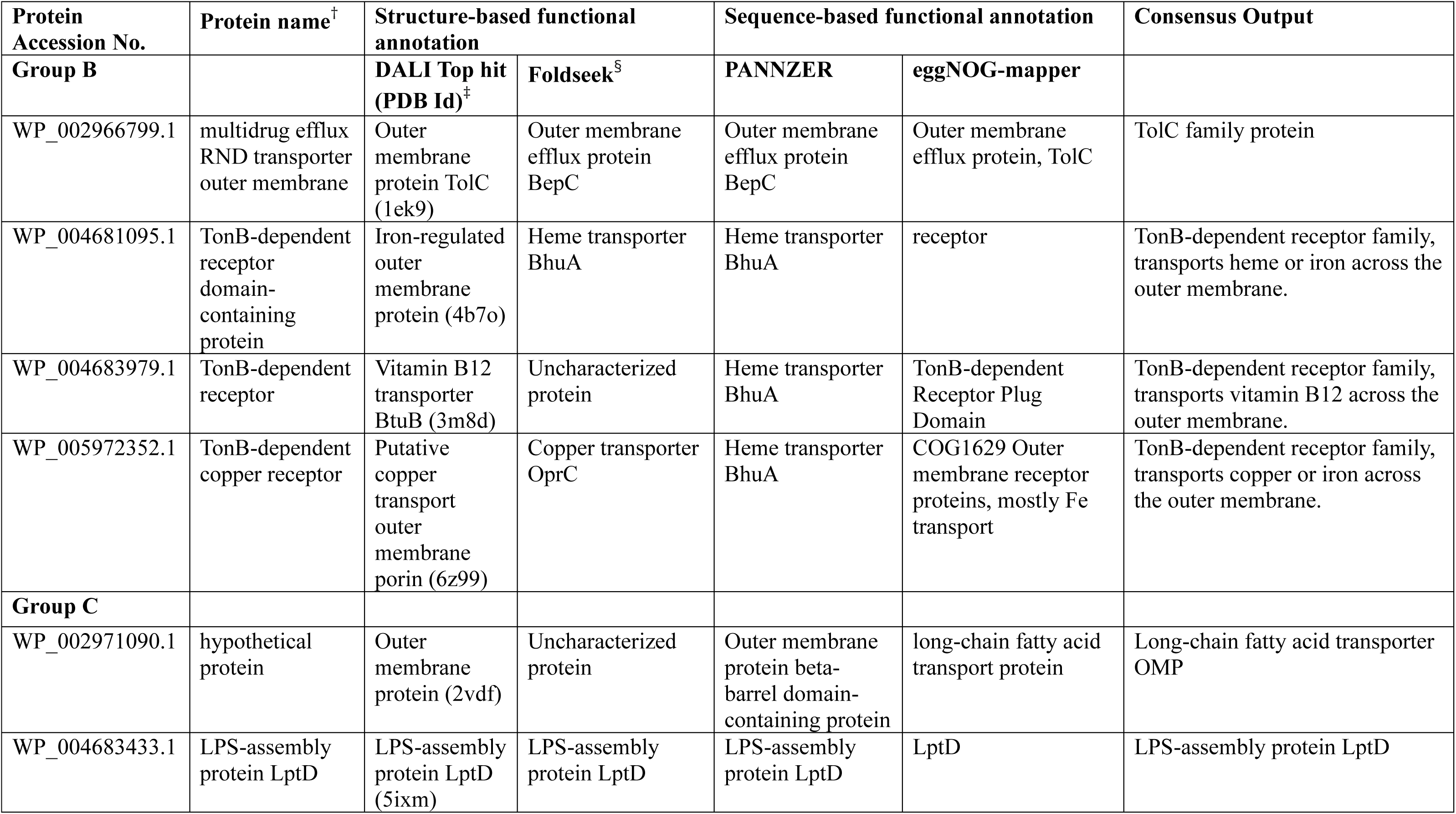
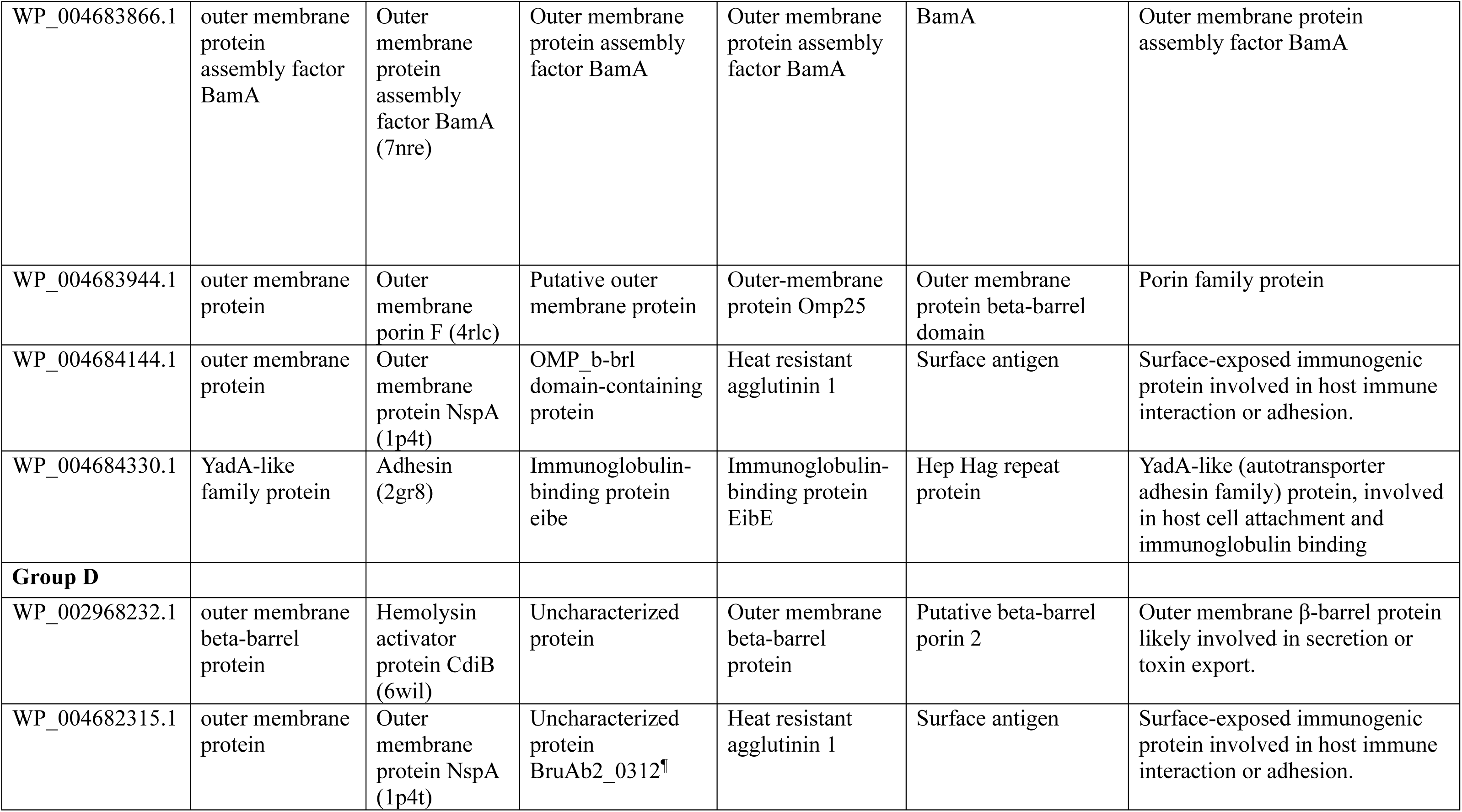
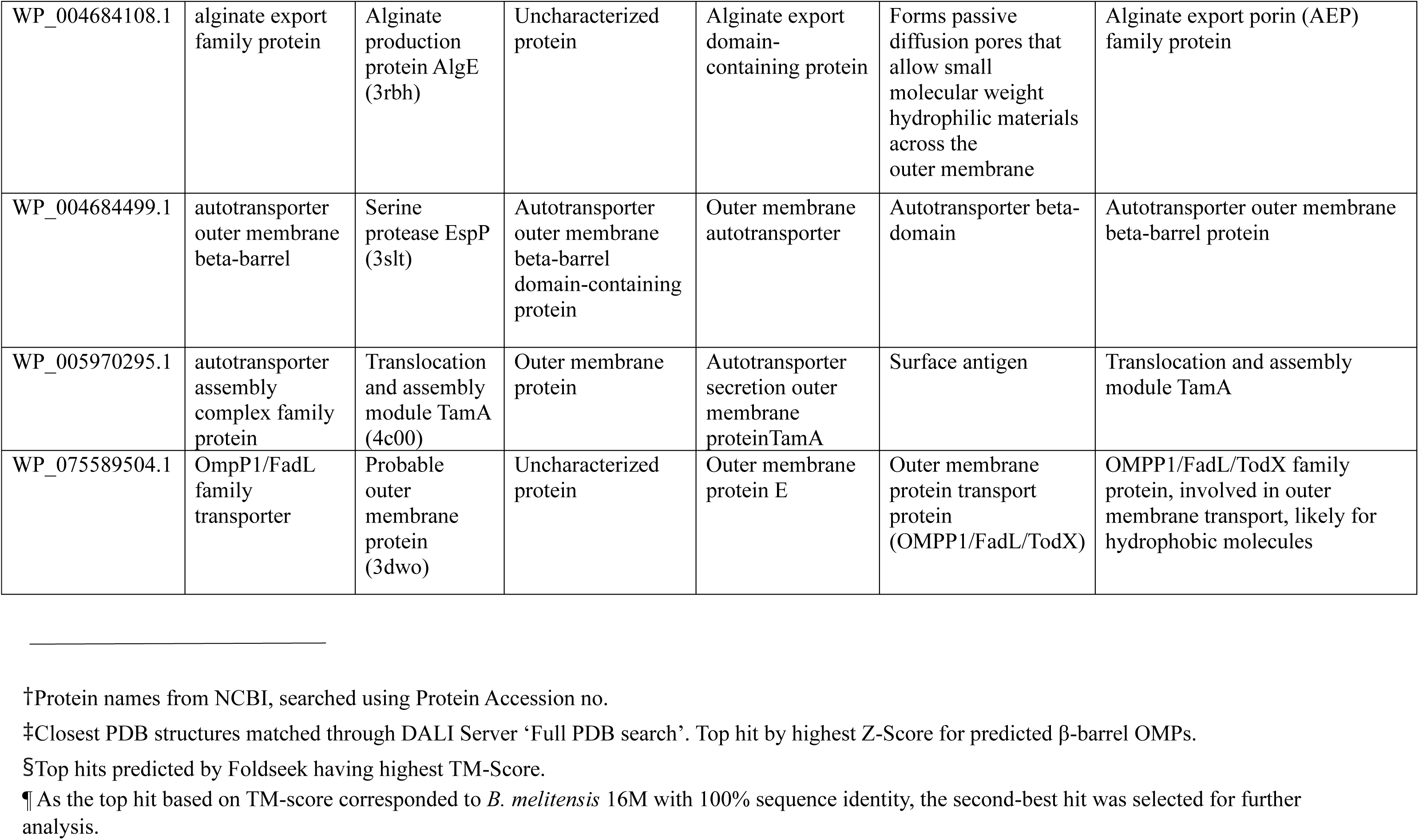
Structure- and sequence-based functional annotation.

| **Protein Accession No.** | **Protein name<sup>†</sup>** | **Structure-based functional annotation** |  | **Sequence-based functional annotation** |  | **Consensus Output** |
| --- | --- | --- | --- | --- | --- | --- |
| **Group B** |  | **DALI Top hit (PDB Id)<sup>‡</sup>** | **Foldseek<sup>§</sup>** | **PANNZER** | **eggNOG-mapper** |  |
| WP_002966799.1 | multidrug efflux RND transporter outer membrane | Outer membrane protein TolC (1ek9) | Outer membrane efflux protein BepC | Outer membrane efflux protein BepC | Outer membrane efflux protein, TolC | TolC family protein |
| WP_004681095.1 | TonB-dependent receptor domain-containing protein | Iron-regulated outer membrane protein (4b7o) | Heme transporter BhuA | Heme transporter BhuA | receptor | TonB-dependent receptor family, transports heme or iron across the outer membrane. |
| WP_004683979.1 | TonB-dependent receptor | Vitamin B12 transporter BtuB (3m8d) | Uncharacterized protein | Heme transporter BhuA | TonB-dependent Receptor Plug Domain | TonB-dependent receptor family, transports vitamin B12 across the outer membrane. |
| WP_005972352.1 | TonB-dependent copper receptor | Putative copper transport outer membrane porin (6z99) | Copper transporter OprC | Heme transporter BhuA | COG1629 Outer membrane receptor proteins, mostly Fe transport | TonB-dependent receptor family, transports copper or iron across the outer membrane. |
| **Group C** |  |  |  |  |  |  |
| WP_002971090.1 | hypothetical protein | Outer membrane protein (2vdf) | Uncharacterized protein | Outer membrane protein beta-barrel domain-containing protein | long-chain fatty acid transport protein | Long-chain fatty acid transporter OMP |
| WP_004683433.1 | LPS-assembly protein LptD | LPS-assembly protein LptD (5ixm) | LPS-assembly protein LptD | LPS-assembly protein LptD | LptD | LPS-assembly protein LptD |
| WP_004683866.1 | outer membrane protein assembly factor BamA | Outer membrane protein assembly factor BamA (7nre) | Outer membrane protein assembly factor BamA | Outer membrane protein assembly factor BamA | BamA | Outer membrane protein assembly factor BamA |
| WP_004683944.1 | outer membrane protein | Outer membrane porin F (4rlc) | Putative outer membrane protein | Outer-membrane protein Omp25 | Outer membrane protein beta-barrel domain | Porin family protein |
| WP_004684144.1 | outer membrane protein | Outer membrane protein NspA (1p4t) | OMP_b-brl domain-containing protein | Heat resistant agglutinin 1 | Surface antigen | Surface-exposed immunogenic protein involved in host immune interaction or adhesion. |
| WP_004684330.1 | YadA-like family protein | Adhesin (2gr8) | Immunoglobulin-binding protein eibe | Immunoglobulin-binding protein EibE | Hep Hag repeat protein | YadA-like (autotransporter adhesin family) protein, involved in host cell attachment and immunoglobulin binding |
| **Group D** |  |  |  |  |  |  |
| WP_002968232.1 | outer membrane beta-barrel protein | Hemolysin activator protein CdiB (6wil) | Uncharacterized protein | Outer membrane beta-barrel protein | Putative beta-barrel porin 2 | Outer membrane β-barrel protein likely involved in secretion or toxin export. |
| WP_004682315.1 | outer membrane protein | Outer membrane protein NspA (1p4t) | Uncharacterized protein BruAb2_0312<sup>¶</sup> | Heat resistant agglutinin 1 | Surface antigen | Surface-exposed immunogenic protein involved in host immune interaction or adhesion. |
| WP_004684108.1 | alginate export family protein | Alginate production protein AlgE (3rbh) | Uncharacterized protein | Alginate export domain-containing protein | Forms passive diffusion pores that allow small molecular weight hydrophilic materials across the outer membrane | Alginate export porin (AEP) family protein |
| WP_004684499.1 | autotransporter outer membrane beta-barrel | Serine protease EspP (3slt) | Autotransporter outer membrane beta-barrel domain-containing protein | Outer membrane autotransporter | Autotransporter beta-domain | Autotransporter outer membrane beta-barrel protein |
| WP_005970295.1 | autotransporter assembly complex family protein | Translocation and assembly module TamA (4c00) | Outer membrane protein | Autotransporter secretion outer membrane protein TamA | Surface antigen | Translocation and assembly module TamA |
| WP_075589504.1 | OmpP1/FadL family transporter | Probable outer membrane protein (3dwo) | Uncharacterized protein | Outer membrane protein E | Outer membrane protein transport protein (OMPP1/FadL/TodX) | OMPP1/FadL/TodX family protein, involved in outer membrane transport, likely for hydrophobic molecules |

†Protein names from NCBI, searched using Protein Accession no.
‡Closest PDB structures matched through DALI Server 'Full PDB search'. Top hit by highest Z-Score for predicted β-barrel OMPs.
§Top hits predicted by Foldseek having highest TM-Score.
¶As the top hit based on TM-score corresponded to *B. melitensis* 16M with 100% sequence identity, the second-best hit was selected for further analysis.

### 2.5 Detection of amino acid sequence variation across 46 *B. melitensis* strains and structural mapping

Predicted OMBB proteins from *B. melitensis* 16M genome were used to search for similar proteins across 46 complete *B. melitensis* genomes using BLASTP (E-value < 1.0E-03, bitscore > 100) to detect sequence variations. Orthologous sequences were aligned using ClustalW for Multiple Sequence Alignment (MSA). Variations identified in the MSA were subsequently mapped onto the structural models in PyMOL (Figures 2, 3).

**Figure 2:**
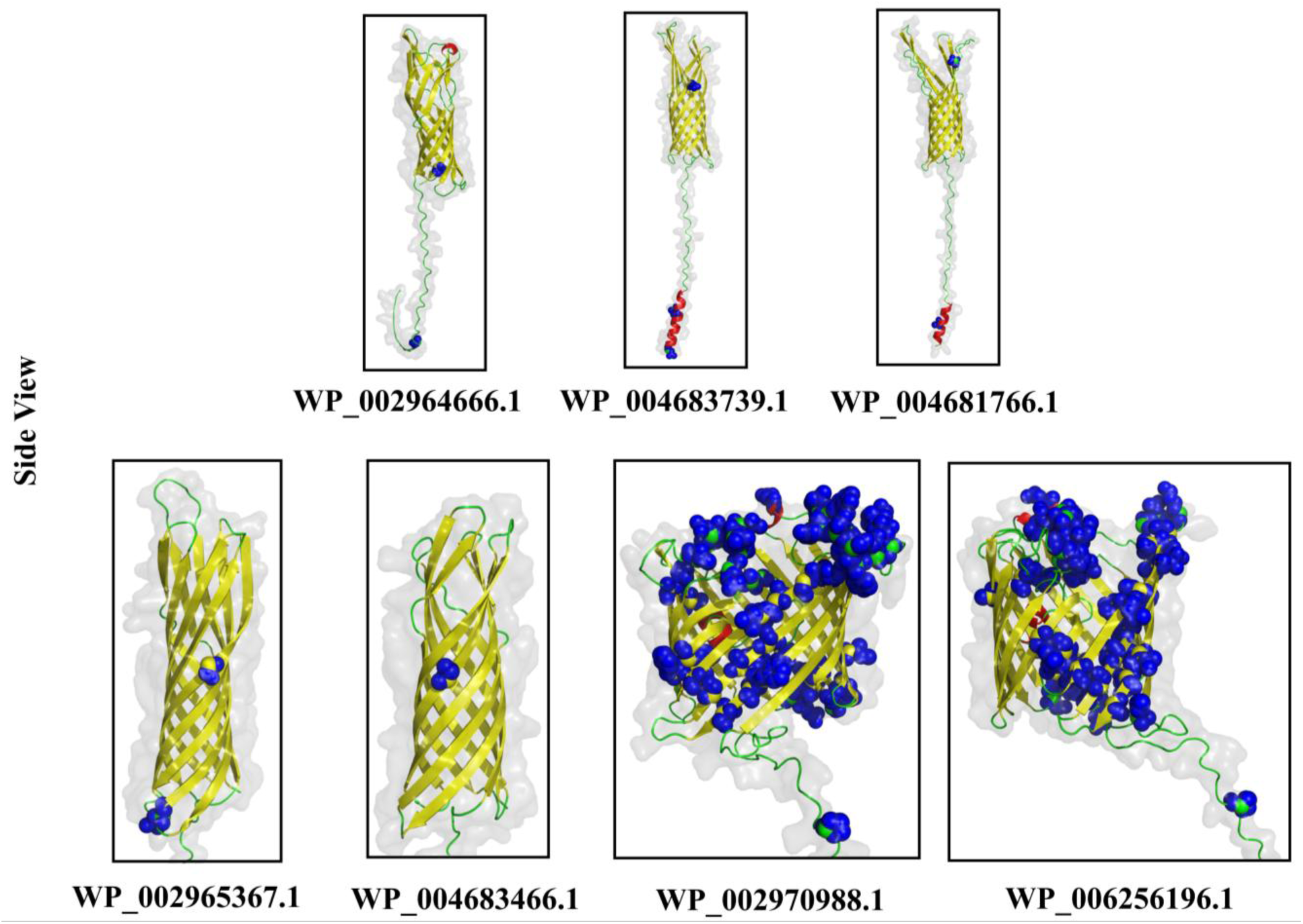
Amino acid sequence variations mapped onto structural models of Group A OMBB proteins. Sequence variation analysis across 46 *B. melitensis* strains and mapping onto the structural models was performed for Group A proteins to determine their genetic diversity. WP_002970988.1 (Omp2a) and WP_006256196.1 (Omp2b) exhibited several variations across entire length of protein. Structures shown here are AlphaFold structures. β-barrel domain, α-helical regions, and loops are shown in yellow, red, and green colours. Blue spheres indicate amino acid variations studied across 46 strains of *B. melitensis*.

**Figure 3:**
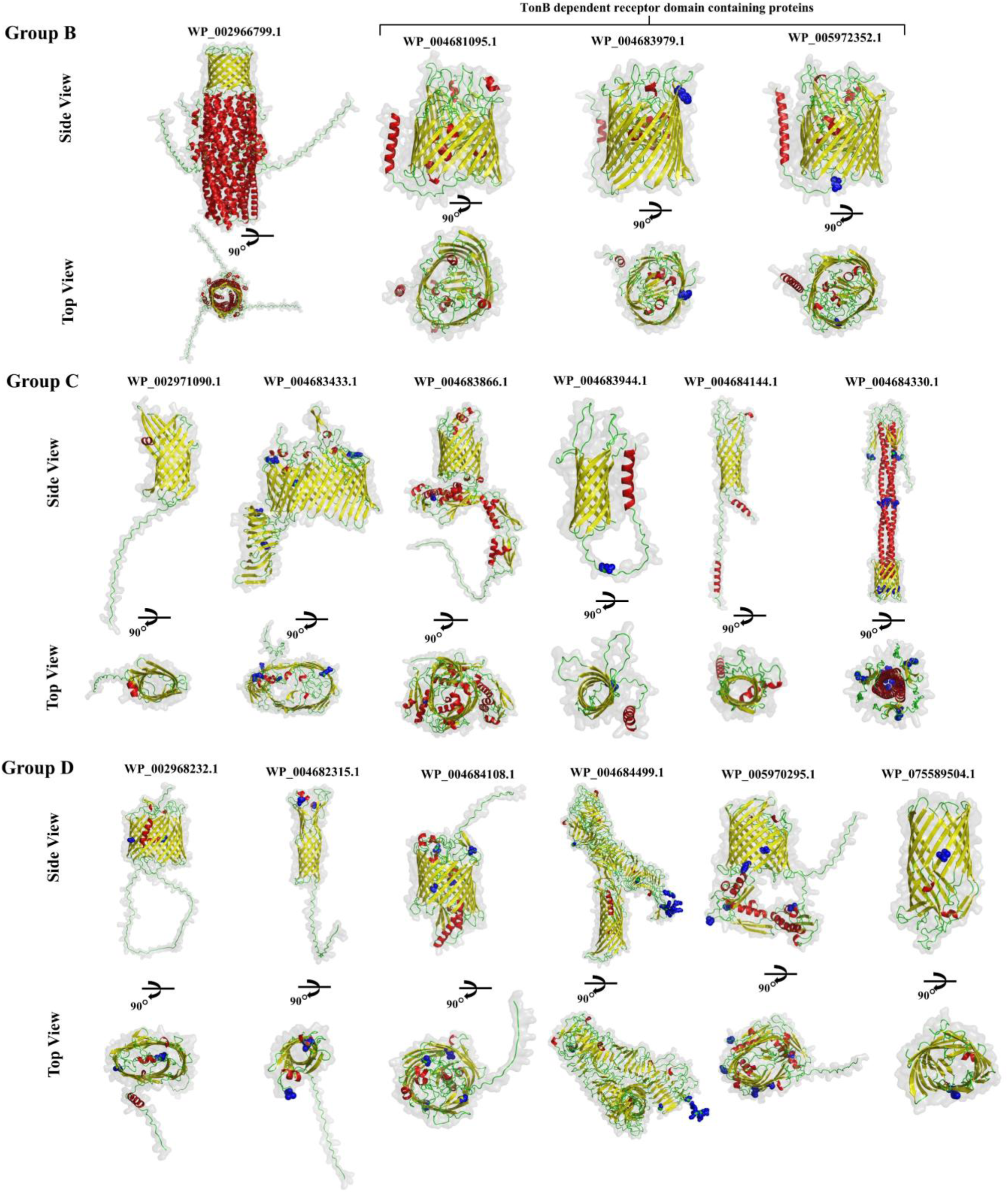
Structural models of proteins in Group B, Group C, and Group D. Group. **B** comprises four proteins: WP_002966799.1, WP_004681095.1, WP_004683979.1 and WP_005972352.1. The trimeric structure of WP_002966799.1 (TolC) consists of three protomers, each with four β-strands creating one-third of the β-barrel. α-helices extend into the periplasm, creating a channel typical of RND-type efflux pumps. WP_004681095.1, WP_004683979.1 and WP_005972352.1 are TonB dependent receptor domain containing proteins. **Group C** comprises WP_002971090.1, WP_004683433.1, WP_004683866.1, WP_004683944.1, WP_004684144.1 and WP_004684330.1. **Group D** comprises six proteins: WP_002968232.1, WP_004682315.1, WP_004684108.1, WP_004684499.1, WP_005970295.1 and WP_075589504.1. β-barrel domain, α-helical regions, and loops are shown in yellow, red, and green colours. Blue spheres indicate amino acid variations studied across 46 *B. melitensis* strains. Structures shown here are AlphaFold 3 structures and β-barrel architecture for all the proteins was validated using ESMFold, SWISS-MODEL, RoseTTAFold, and TrRosetta.

### 2.6 Structural prediction of protein complexes

To identify the components of the RND efflux pumps, Lpt, and Bam complexes, the proteome of *B. melitensis* 16M was manually searched. FASTA sequence for each annotated protein of the complex was retrieved from NCBI. Complex structures were modelled with AlphaFold3, and protein-protein interactions (PPIs) were analyzed using PDBePISA.^51^

#### 2.6.1 RND (Resistance-Nodulation-Division) efflux pumps

In 2024, H. Ayoub et al., identified 5 efflux related proteins-BepC, BepD, BepE, BepF, and BepG- in *B. melitensis*.^52^ BLASTP analysis showed that BepC (WP_002966799.1) is homologous to TolC family protein. Similarly, BepD (WP_004685422.1) and BepF (WP_004681660.1) are homologs of RND transporter periplasmic adaptor subunit AcrA. BepE (WP_004686764.1) and BepG (WP_004686857.1) are homologous to efflux RND transporter permease subunit. Oligomeric states were modelled with AlphaFold 3: a trimer for WP_002966799.1, hexamers for WP_004685422.1 and WP_004681660.1, and trimers for WP_004686764.1 and WP_004686857.1. The architecture of AcrAB-TolC (PDB Id: 5NG5) was used as a template to assemble RND efflux pumps, assigning WP_002966799.1 as Outer Membrane Factor (OMF), WP_004685422.1 and WP_004681660.1 as Membrane Fusion Proteins (MFPs), and WP_004686764.1 and WP_004686857.1 as Inner Membrane Proteins (IMPs). PPIs were evaluated using PDBePISA: monomers of BepC and BepE were analyzed against the subunits of BepD, and monomers BepC and BepG against the subunits of BepF. Interactions were visualized in PyMOL (Figure 4).

**Figure 4:**
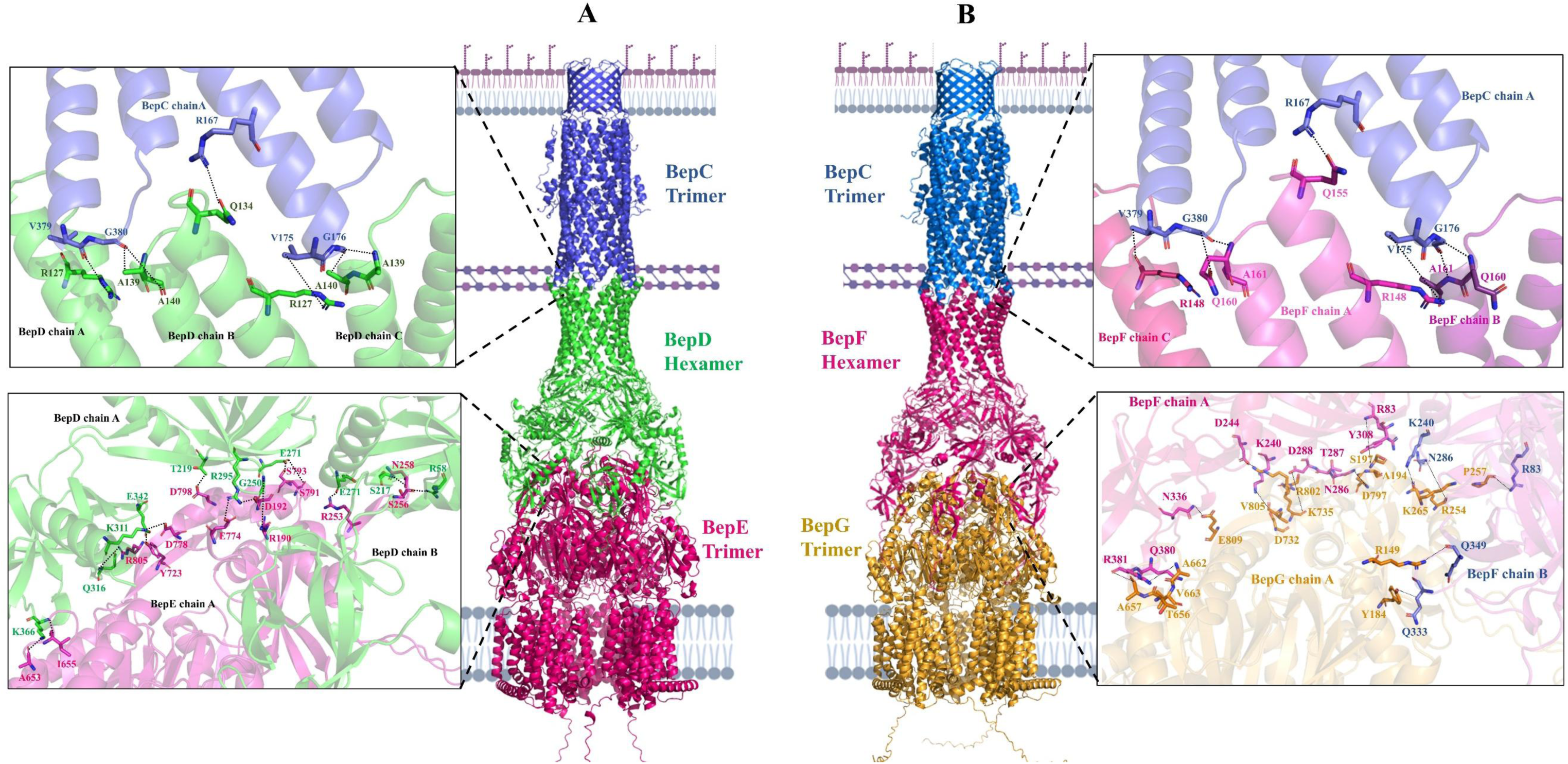
Structural models of *B. melitensis* RND efflux pumps. RND efflux pump forms a tripartite complex consisting of two trimeric proteins, inner membrane proteins (IMP) and outer membrane factors (OMF), bridged by hexameric membrane fusion proteins (MFP). Homology based study have predicted that BepC (WP_002966799.1) classified in Group B is a trimeric OMF in *B. melitensis*. **A. BepDE-BepC efflux pump:** BepC is a TolC homolog that is associated with BepD (WP_004685422.1). BepD, a hexameric membrane fusion protein (MFP), functions as a periplasmic adaptor that further interacts with BepE (WP_004686764.1). BepE is an RND transporter permease located in the inner membrane, involved in the translocation of a wide range of molecules across the bacterial membrane. Enlarged view of PPIs represents the hydrogen bonds between monomers of each BepC and BepE with subunits of periplasmic BepD. **B. BepFG-BepC efflux pump:** BepC is also associated with other RND efflux pump having BepF (WP_004681660.1) as MFP and BepG (WP_004686857.1) as IMP. Enlarged view shows hydrogen bonds between monomers of each BepC and BepG with subunits of BepF.

#### 2.6.2 Lpt complex

The Lpt complex is well characterized in *E. coli*, comprising seven essential proteins (LptA–G).^53^ The complete proteome of *B. melitensis* available in the NCBI database was manually searched to identify annotated homologs of these components in *B. melitensis*. The complex comprises two subcomplexes linked by the periplasmic lipoprotein LptA (WP_004686649.1). The outer-membrane subcomplex contains LptD (WP_004683433.1) and LptE (WP_002964883.1); the inner-membrane subcomplex contains LptC (WP_004684681.1), LptF (WP_004683435.1), LptG (WP_002967526.1), and the dimeric ATPase LptB (WP_002971717.1). Structures of LptDE and LptB_2_FGC were predicted using AlphaFold 3. The LptB_2_FGC model was aligned to the *Vibrio cholerae* LptB_2_FGC crystal structure (PDB Id: 6MJP) using US-align, which yielded an RMSD of 6.69 Å. Structure of LptA was generated separately using AlphaFold 3. These structures were then manually assembled to form a complete Lpt complex in *B. melitensis.* Protein–protein interfaces between β-jelly rolls of LptD-LptA and LptA- LptC were analyzed in PDBePISA, and hydrogen bonds were visualized in PyMOL (Figure 5).

**Figure 5:**
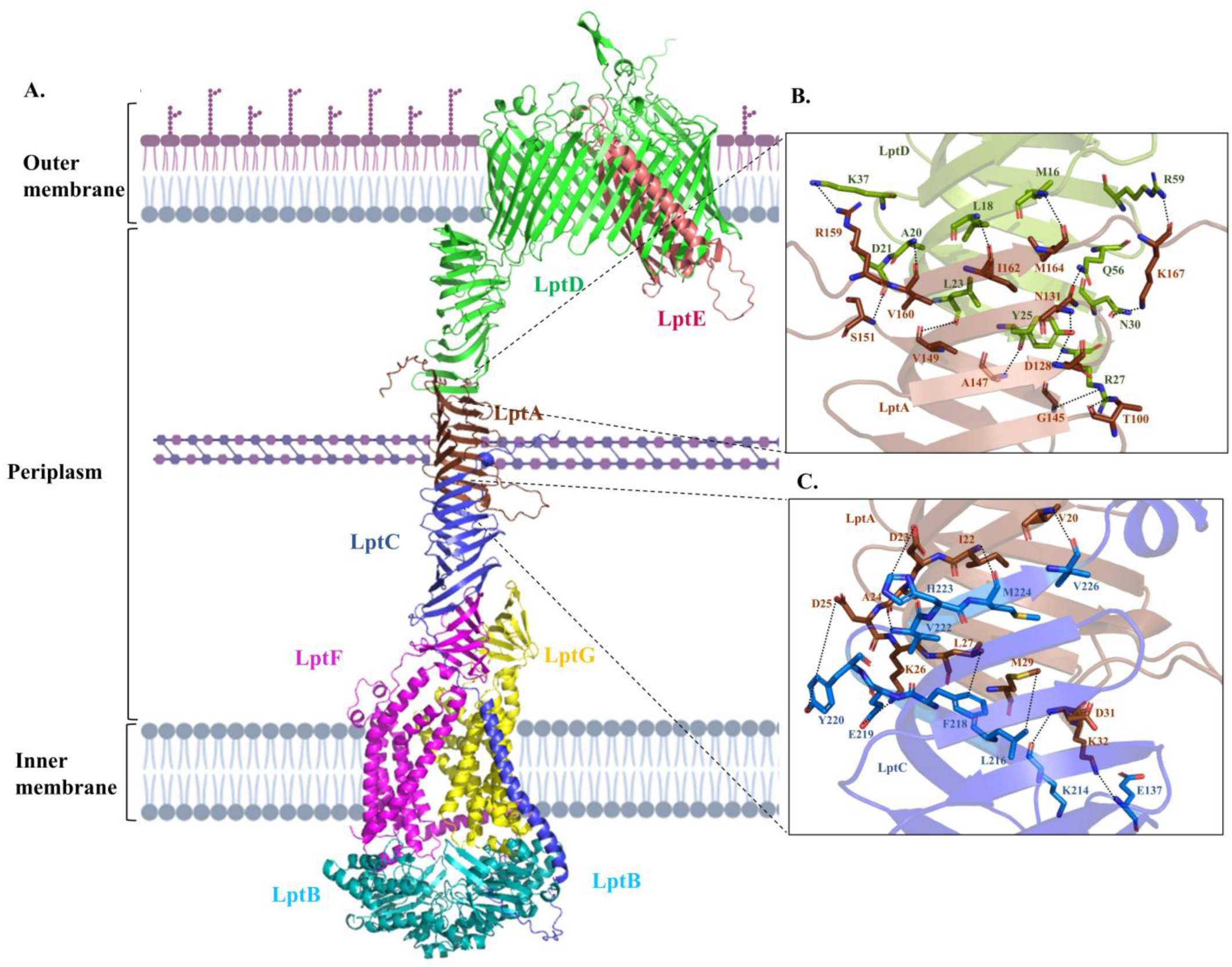
Structural Model of *B. melitensis* Lpt Complex. Lpt machinery consists of two sub-assemblies: LptDE and LptB_2_FG. At the outer membrane, LptD (WP_004683433.1), a 26 stranded β-barrel protein, forms a complex with lipoprotein LptE (WP_002964883.1). LptE functions as a plug that occupies the lumen of LptD and regulates its opening. At the inner membrane, subcomplex LptB_2_FG consists of transmembrane polytopic proteins LptF (WP_004683435.1) and LptG (WP_002967526.1) along with an ABC transporter which comprises of dimeric LptB (WP_002971717.1) towards cytoplasm. LptB_2_FG associate with LptC (WP_004684681.1) which is an auxiliary inner membrane protein. The two subcomplexes are connected by a periplasmic lipoprotein LptA (WP_004686649.1).

#### 2.6.3 BAM complex

We modelled the BAM complex using AlphaFold 3 with sequences of BamA (WP_004683866.1), BamD (WP_002964530.1), and BamE (WP_005969315.1), as BamB and BamC are absent in *B. melitensis.*^54^ Structure-based sequence alignment of *B. melitensis* BamA with *E. coli* BamA (UniProt Id: P0A940) was performed with PROMALS3D^55^ to identify equivalent interface residues. Protein– protein interface residues among BamA, BamD and BamE were further predicted using PDBePISA (Figure 6).

**Figure 6:**
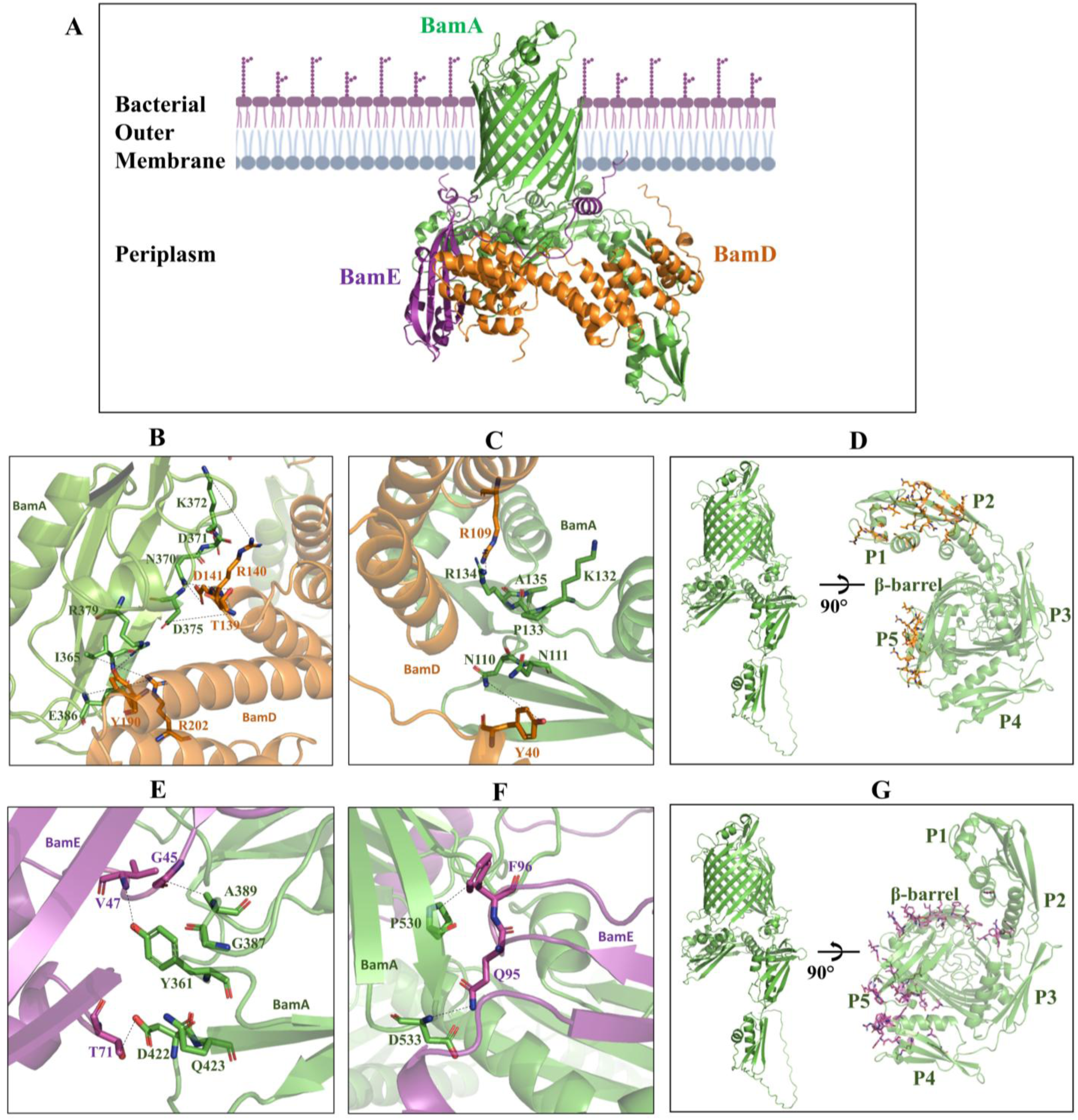
The Bam Complex of *B. melitensis*. **(A)** The **Bam complex** in *B. melitensis* consists of BamA, BamD, and BamE. BamA is a 16-stranded outer membrane β-barrel (OMBB) protein with five POTRA domains at its N-terminal towards periplasm, which facilitate interactions with the periplasmic lipoproteins BamD and BamE. Structure-based sequence alignment of *B. melitensis* BamA (BamA^Bm^) with *E. coli* BamA (BamA^Ec^) revealed key interaction sites. **BamA^Bm^-BamD^Bm^ interactions: (B)** Highlights the equivalent residues in POTRA5 of BamA^Bm^ that interact with BamD^Bm^. **(C)** Displays the corresponding residues in POTRA2 of BamA^Bm^ involved in interactions with BamD^Bm^. **(D)** Represents additional interfacing residues in BamA^Bm^ predicted by PDBePISA involved in interaction with BamD^Bm^. **BamA^Bm^-BamE^Bm^ interactions: (E)** Shows the equivalent residues in POTRA5 of BamA^Bm^ interacting with BamE^Bm^. **(F)** Highlights residues in periplasmic loop 3 of BamA^Bm^ involved in interaction with BamE^Bm^. **(G)** Depicts additional residues of BamA^Bm^ predicted by PDBePISA that interact with BamE^Bm^. Dotted lines represent hydrogen bonds between amino acid residues.

## 3. Results and Discussion

### 3.1 Prediction of OM β-barrel proteins

*B. melitensis* 16M (genome assembly ASM74041v1) encodes 2978 proteins. We applied a consensus-based computational pipeline to identify OM β-barrel proteins from this proteome. Nine tools were used for proteome analysis: Pepstats, CELLO, PSORTb, MCMBB, OMPdb, TMbed, TMBETADISC- RBF, SignalP, and SPAAN (Table S1). For OMBB protein prediction specifically, we considered four OMBB prediction tools (MCMBB, OMPdb, TMbed, and TMBETADISC-RBF). 24 OMBB proteins were predicted positive by all four tools and were considered for further analysis (Table S1). These proteins were grouped based on the extent of prior work in *B. melitensis* determined through literature survey. Group A (eight proteins) have experimental OMP evidence and in silico structural models (major OMPs: Omp2a, Omp2b, Omp25, Omp31); Group B (four proteins) have experimental OMP evidence but lack structural predictions; Group C (six proteins) have only in silico studies; Group D (six proteins) comprises novel β-barrel proteins identified in this study, merely annotated in NCBI and/or UniProt with no prior experimental evidence or computational characterization. (Tables 1, 2).

### 3.2 OMBB protein validation of uncharacterized proteins

Because the 12 proteins in Groups C and D have no experimental data, we first had to confirm that they are OMBB candidates rather than false positives from the discovery pipeline. We therefore applied four complementary tools-Protein GRAVY (hydrophobicity), PPM 3.0 (membrane orientation and ΔG_transfer), DREAMM (residue-level membrane contacts), and MemProtMD_Insane (coarse-grained MD stability)- to test whether each protein could stably embed in the outer membrane. Together, these independent tools provided convergent evidence for outer-membrane insertion. Protein GRAVY (Grand Average of Hydropathy) calculates the mean hydropathy of a protein sequence, indicating its hydrophobicity (water-repelling) or hydrophilicity (water-attracting) nature. Values ranging between −1.5 to 0.5 typically indicate hydrophobic character. All 12 proteins had GRAVY scores from −0.504 to −0.09, consistent with membrane association. PPM 3.0 estimates optimal protein positioning in lipid bilayers and computes transfer free energy (ΔG; typically −400 to −10 kcal/mole) for moving a protein from water into a membrane-like environment. Group C and D proteins generated values within this range, showing high probability of being embedded in transmembrane region. Transmembrane segments identified by PPM 3.0 embedded in the membrane are shown in Table 5. Given the GRAVY and PPM 3.0 results, we identified membrane-penetrating residues (Table 5) using DREAMM, further validating OM insertion. Additionally, MemProtMD_Insane was employed to assess the stability of the protein within a membrane environment. MemProtMD_Insane embedded each protein into lipid bilayer and simulated its dynamics using coarse-grained molecular dynamics. The aligned RMSD remained consistently low throughout the simulation, indicating stable protein conformations in the lipid membrane.

**Table 5:**
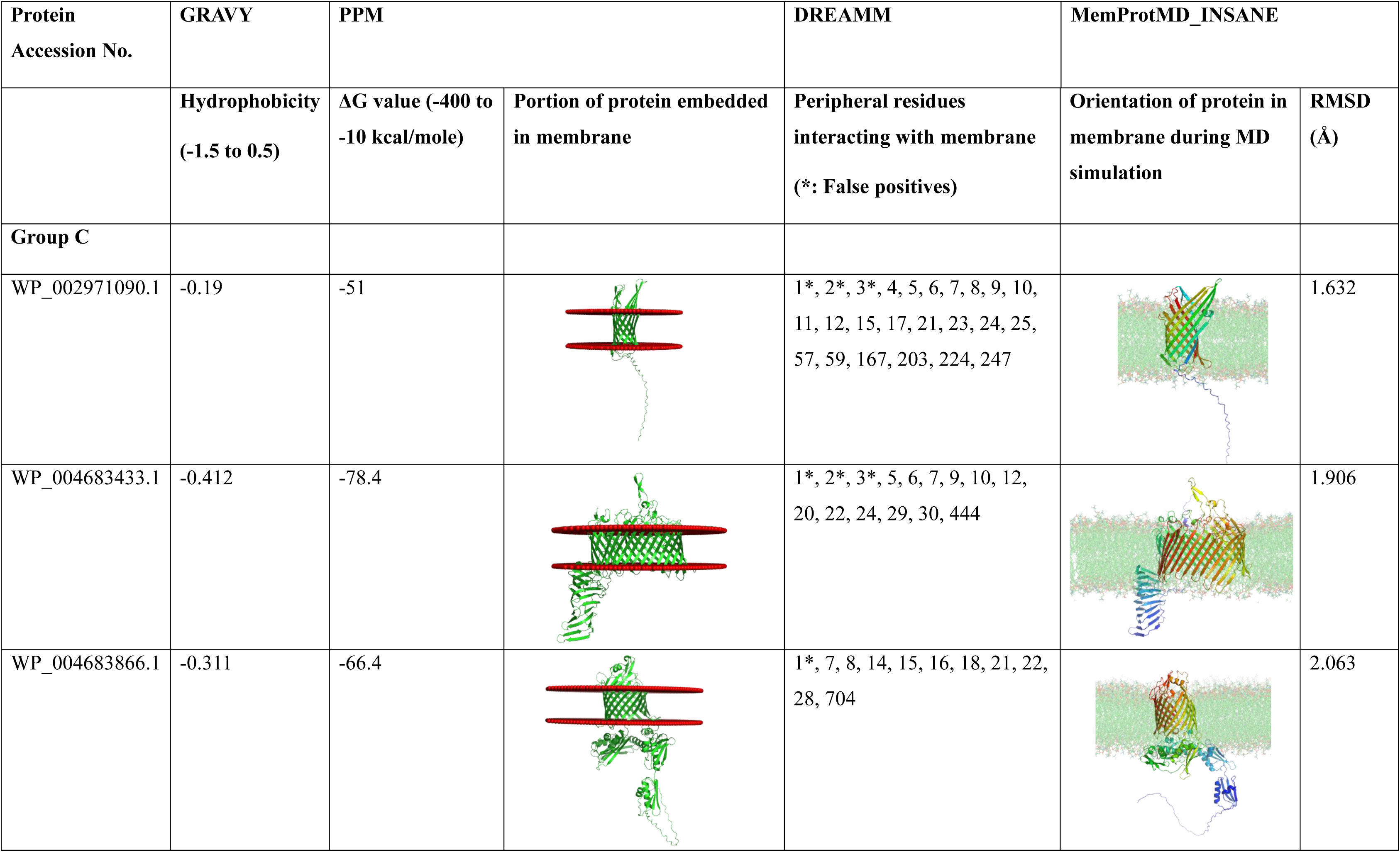
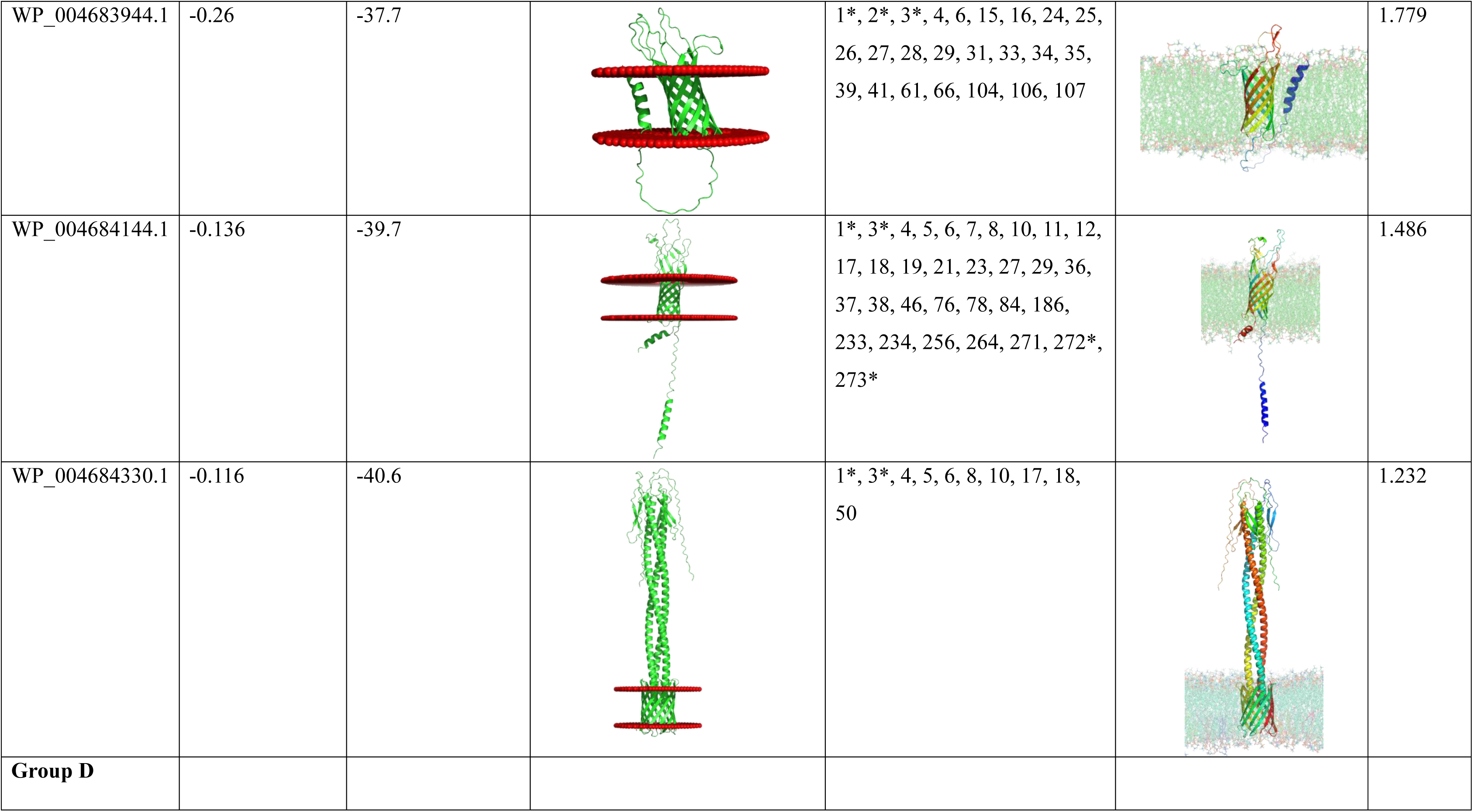
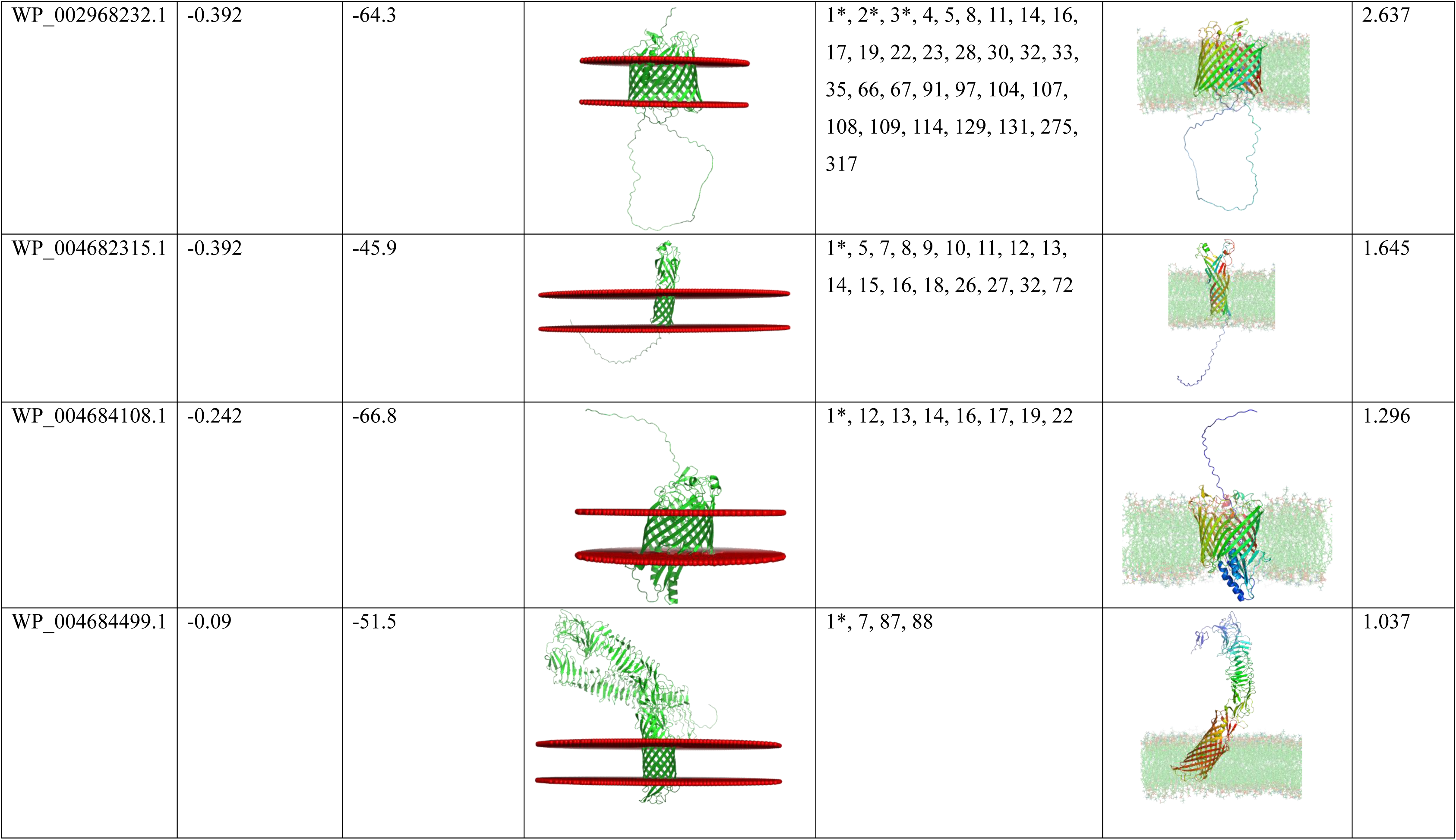
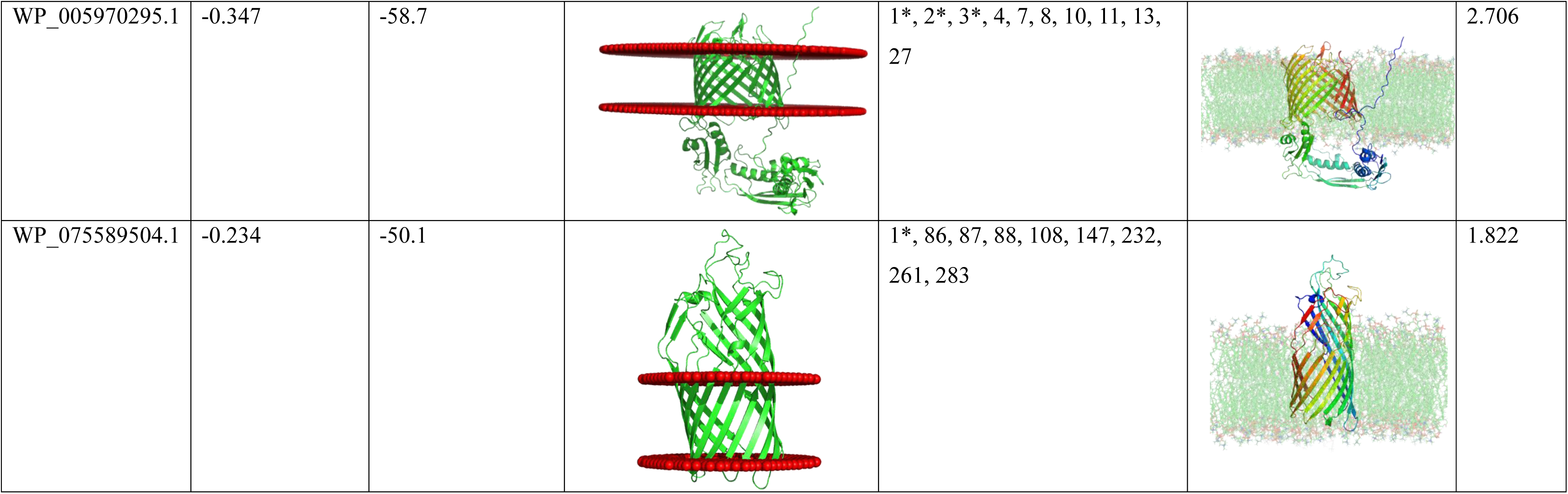
Validation of Outer membrane proteins insertion in lipid membrane.

| Protein<br>Accession No. | GRAVY | PPM |  | DREAMM | MemProtMD_INSANE |  |
| --- | --- | --- | --- | --- | --- | --- |
| | Hydrophobicity<br>(-1.5 to 0.5) | $\Delta G$ value (-400 to<br>-10 kcal/mole) | Portion of protein embedded<br>in membrane | Peripheral residues<br>interacting with membrane<br>(*: False positives) | Orientation of protein in<br>membrane during MD<br>simulation | RMSD<br>(Å) |
| **Group C** |  |  |  |  |  |  |
| WP_002971090.1 | -0.19 | -51 |  | 1*, 2*, 3*, 4, 5, 6, 7, 8, 9, 10,<br>11, 12, 15, 17, 21, 23, 24, 25,<br>57, 59, 167, 203, 224, 247 |  | 1.632 |
| WP_004683433.1 | -0.412 | -78.4 |  | 1*, 2*, 3*, 5, 6, 7, 9, 10, 12,<br>20, 22, 24, 29, 30, 444 |  | 1.906 |
| WP_004683866.1 | -0.311 | -66.4 |  | 1*, 7, 8, 14, 15, 16, 18, 21, 22,<br>28, 704 |  | 2.063 |
| WP_004683944.1 | -0.26 | -37.7 |  | 1*, 2*, 3*, 4, 6, 15, 16, 24, 25, 26, 27, 28, 29, 31, 33, 34, 35, 39, 41, 61, 66, 104, 106, 107 |  | 1.779 |
| WP_004684144.1 | -0.136 | -39.7 |  | 1*, 3*, 4, 5, 6, 7, 8, 10, 11, 12, 17, 18, 19, 21, 23, 27, 29, 36, 37, 38, 46, 76, 78, 84, 186, 233, 234, 256, 264, 271, 272*, 273* |  | 1.486 |
| WP_004684330.1 | -0.116 | -40.6 |  | 1*, 3*, 4, 5, 6, 8, 10, 17, 18, 50 |  | 1.232 |
| **Group D** |  |  |  |  |  |  |
| WP_002968232.1 | -0.392 | -64.3 |  | 1*, 2*, 3*, 4, 5, 8, 11, 14, 16, 17, 19, 22, 23, 28, 30, 32, 33, 35, 66, 67, 91, 97, 104, 107, 108, 109, 114, 129, 131, 275, 317 |  | 2.637 |
| WP_004682315.1 | -0.392 | -45.9 |  | 1*, 5, 7, 8, 9, 10, 11, 12, 13, 14, 15, 16, 18, 26, 27, 32, 72 |  | 1.645 |
| WP_004684108.1 | -0.242 | -66.8 |  | 1*, 12, 13, 14, 16, 17, 19, 22 |  | 1.296 |
| WP_004684499.1 | -0.09 | -51.5 |  | 1*, 7, 87, 88 |  | 1.037 |
| WP_005970295.1 | -0.347 | -58.7 |  | 1*, 2*, 3*, 4, 7, 8, 10, 11, 13, 27 |  | 2.706 |
| WP_075589504.1 | -0.234 | -50.1 |  | 1*, 86, 87, 88, 108, 147, 232, 261, 283 |  | 1.822 |

### 3.3 Identification anxpd characterization of OM Proteins

#### 3.3.1 Group A

Group A comprises eight proteins– WP_002964666.1, WP_002965367.1, WP_002965368.1, WP_004683466.1, WP_004683739.1, WP_002970988.1, WP_004681766.1, and WP_006256196.1– previously identified and characterized as OMBB proteins in *B. melitensis*. Prediction of these proteins through our findings validates the robustness of the pipeline. This group includes an OmpW family protein, Omp2a, Omp2b, Omp31, and four proteins from Omp25 family. Although these proteins have been extensively studied experimentally and computationally, no crystal structures are available. Sequence variations were analyzed across 46 strains of *B. melitensis* and mapped onto the structural models generated with AlphaFold 3 (Figure 2). Most of the proteins were highly conserved; however, Omp2a and Omp2b exhibited greater amino acid variability, suggesting functional divergence or selective pressure driven by host immune recognition. Because these OMBB proteins are already well-characterized, they were not analyzed further in this study.

#### 3.3.2 Group B

Group B contains proteins that have been studied experimentally but lack predicted structures: WP_002966799.1, WP_004681095.1, WP_004683979.1, and WP_005972352.1 (Table 2).

##### 3.3.2.1 WP_002966799.1 TolC family protein

WP_002966799.1 is annotated in NCBI as multidrug efflux RND transporter BepC and in UniProt as a homolog of TolC family protein. TolC is a trimeric 12-stranded α/β-barrel, spanning the outer membrane of Gram-negative bacteria and is a key component of efflux systems such as AcrAB−TolC in Enterobacteriaceae and MexAB−OprM, MexCD−OprJ, and MexXY−OprM in *Pseudomonas aeruginosa.*^56,57^ Together with an inner membrane protein (IMP), a membrane fusion protein (MFP), TolC-like proteins form the tripartite Resistance-Nodulation-Division (RND) pump that expels toxic compounds across the cell envelope.^58^ BepC is a homotrimer with each protomer comprising four transmembrane strands (Figure 3). The consensus results from structural alignment tools (DALI and Foldseek) and sequence annotation tools (PANNZER and eggNOG-mapper) supported classification of BepC as a TolC-family multidrug efflux RND transporter (Table 4). Sequence variation analysis showed that BepC is a highly conserved protein, with no amino acid variation detected (Figure 3, Table S3).

##### 3.3.2.2 TonB-dependent receptor domain-containing proteins

We identified three TonB-dependent proteins (TBDP): WP_004681095.1, WP_004683979.1, and WP_005972352.1. Each protein forms a 22-stranded β-barrel (Figure 3). Danese et al. in 2004, identified three TonB iron receptors encoding genes (BMEII0105, BMEII0297, and BMEI0657), and showed that the TonB system contributes to 2,3-dihydroxybenzoic acid (DHBA) utilization and iron assimilation in *B. melitensis.*^59^ WP_004681095.1 and WP_005972352.1 are annotated as Heme transporter BhuA in UniProt. BhuA has been identified as a potential vaccine candidate in *B. melitensis* 16M.^60^ WP_004681095.1 showed the highest Z-score match with iron-regulated outer membrane protein FrpB (PDB Id: 4B7O) of *Neisseria meningitidis* (Table 4). WP_005972352.1 showed the highest DALI Z-score match to a putative copper transport porin OprC (PDB Id: 6Z99) of *P. aeruginosa* (Z-score = 57.9, RMSD = 1 Å). Foldseek analysis predicted WP_004681095.1 and WP_005972352.1 homologous to heme transporter BhuA and copper transporter OprC, respectively. These findings suggest that *B. melitensis* possesses distinct outer membrane proteins specialized for the uptake of essential metals such as iron and copper. WP_004683979.1, aligned (Z score = 45, RMSD = 2Å) with the vitamin B12 transporter BtuB (PDB Id: 3M8D) from *E. coli* in DALI, whereas Foldseek aligned it with an uncharacterized protein. Sequence-based annotation tools assigned broader groups: PANNZER identified all three proteins as BhuA, while eggNOG-mapper placed them in membrane-receptor orthologous groups. All three possess a large N-terminal plug domain that likely regulates transport across the cell envelope. US-align superpositions indicated overall structural similarity among the three proteins (RMSD=3.62Å). Sequence variation analysis across 46 *B. melitensis* strains showed high conservation. A single variation was detected in WP_004683979.1 (W315) and WP_005972352.1 (N358) (Figure 3, Table S3).

#### 3.3.3 Group C

Proteins in Group C have been predicted and studied in silico in *B. melitensis*, but they have not yet been examined experimentally. These proteins include: WP_002971090.1, WP_004683433.1, WP_004683866.1, WP_004683944.1, WP_004684144.1, and WP_004684330.1 (Table 2).

##### 3.3.3.1 WP_002971090.1

WP_002971090.1 is annotated in NCBI as hypothetical protein and in UniProt as outer membrane β-barrel protein. WP_002971090.1 has been predicted as a cross-protective potential antigen against pathogenic *Brucella* spp. and can act as an adhesin.^23^ Structural model generated by five tools revealed a ten-stranded β-barrel architecture (Figure 3). Four of the β-strands are extended towards the extracellular side, which might be involved in interactions with the host environment. WP_002971090.1 was predicted to have adhesin-like properties by SPAAN (P_ad_ value = 0.73). It showed the best structural alignment, in DALI, with the outer membrane protein OpcA (PDB Id: 2VDF) of *N. meningitidis* (Z-score = 17, RMSD = 3.5 Å) (Table 4). OpcA acts as an adhesin that binds to sialic acid-containing polysaccharides on the surface of epithelial cells.^61^ Foldseek predicted homology to an uncharacterized OMP of *Yersinia bercovieri*. Sequence-based annotation classified it as an OMBB orthologous to a long-chain fatty acid transport protein. Combined results from all the annotation tools indicate that WP_002971090.1 likely functions as an adhesin and a transporter of bulky substrates (polysaccharide or long chain fatty acids). Sequence variation analysis of the protein across 46 *B. melitensis* strains revealed that it is a conserved protein with no variation (Figure 3, Table S3).

##### 3.3.3.2 WP_004683433.1 LptD

This protein is annotated in NCBI and UniProt as LptD. LptD is the outer membrane component of LPS transport complex. It is a 26-stranded β-barrel protein having a distinctive periplasmic domain comprising a β-jelly roll domain (Figure 3). Servais et al. in 2023 predicted the structure of *B. abortus* LptD and observed its localization in the outer membrane.^62^ In *B. melitensis*, LptD has been computationally identified as a potential vaccine candidate against brucellosis.^60,63^ SPAAN analysis predicted WP_004683433.1 as a potential adhesin (P_ad_ value=0.64). All four functional annotation tools (DALI, Foldseek, PANNZER, and eggNOG-mapper) predicted it as a homolog of the LPS assembly protein LptD (Table 4). Multiple Sequence Alignment of WP_004683433.1 homologs across 46 *B. melitensis* strains revealed five variations present in β-jelly roll domain (Q45, E215), and extracellular loop (ECL) region (Q292, G347, and R554) (Figure 3, Table S3).

##### 3.3.3.3 WP_004683866.1 BamA

WP_004683866.1 is annotated as outer membrane protein assembly factor BamA in NCBI and UniProt. Structure of WP_004683866.1 constructed using five modelling tools predicted a 16-stranded β-barrel architecture and five polypeptide transport-associated (POTRA) domains (Figure 3). BamA is a key component of *<u>β</u>*-barrel <u>a</u>ssembly <u>m</u>achine (BAM) complex that mediates folding and insertion of OMPs in outer membrane of Gram-negative bacteria. It has a β-barrel domain at C-terminal in OM, and five POTRA domains at N-terminal in periplasm. The barrel exhibits a lateral opening between strands β1 and β16 for insertion of OMPs in OM.^64^ In 2008, Gatsos X et al. conducted a study in *B. melitensis* to find components of OM assembly pathway using Hidden Markov Models (HMMs). They found that the genus *Brucella* encodes homologs of BamA, BamD, and BamE proteins but lacks BamB and BamC proteins as present in *E. coli* and other Gamma-proteobacteria.^18,65^ The structure-based functional annotation using DALI server revealed the closest match (Z-score = 33.1 and RMSD = 2.5 Å) with BamA of *E. coli* O157:H7 (PDB Id: 7NRE) (Table 4). Foldseek analysis also aligned this protein with homologous BamA protein of *Ensifer adhaerens* OV14, having highest TM score. Sequence-based annotation using PANNZER and eggNOG-mapper described it as BamA with PPV value of 0.6. Out of the 46 *B. melitensis* strains included in the sequence variation study, one strain (*B. melitensis* QY1) lacked the BamA protein in its annotated proteome and was therefore excluded from the analysis. Consequently, only 45 strains were used for amino acid variation analysis. Sequence comparison of WP_004683866.1 across 45 strains of *B. melitensis* revealed only one variation at V318 in POTRA 4 domain (Figure 3, Table S3).

##### 3.3.3.4 WP_004683944.1

WP_004683944.1 is annotated as OMP in NCBI and a β-barrel domain-containing protein in UniProt. All five structural modeling tools revealed a β-barrel architecture comprising eight anti-parallel β-strands (Figure 3). A previous in silico study has predicted this protein as a cross-protective potential antigen against pathogenic *Brucella* spp.^23^ In our study, SPAAN predicted WP_004683944.1 to be an adhesin (P_ad_ value = 0.82). DALI search showed the best structural alignment with *P. aeruginosa* OprF (PDB Id: 4RLC) (Z-score = 16.4, RMSD = 2.1 Å; Table 4), a general porin that facilitates the non-specific diffusion of ionic species and small polar nutrients, including polysaccharides. It also anchors the OM to the peptidoglycan layer, contributes in adhesion to eukaryotic cells and biofilm formation under anaerobic conditions.^66^ Structural similarity with OprF suggested that WP_004683944.1 may have similar functions. Foldseek has annotated this protein as an OMBB. Sequence-based annotation using PANNZER has predicted it as Omp25, which is a well characterized protein in *Brucella* spp., while eggNOG-mapper found it as an ortholog of OMBB protein. These consistent results strongly suggest that the protein likely belongs to the Porin family, which plays a crucial role in nutrient transport and host-pathogen interactions in Gram-negative bacteria. This also suggests that WP_004683944.1 may have evolved from Omp25 family proteins having a common ancestor. Multiple Sequence Alignment of WP_004683944.1 homologs across 46 *B. melitensis* strains revealed only one variation in intracellular loop (ICL) region (N36), indicating that it is a highly conserved protein (Figure 3, Table S3).

##### 3.3.3.5 WP_004684144.1

WP_004684144.1 is annotated as OMP in NCBI and OMBB protein in UniProt. Structural model of WP_004684144.1 predicted with five tools revealed a β-barrel architecture consisting of eight anti-parallel β-strands (Figure 3). WP_004684144.1 has been predicted as a cross-protective potential antigen against pathogenic *Brucella* spp. by Hisham et al. in 2018.^23^ WP_004684144.1 showed best match with outer membrane protein NspA (PDB Id: 1P4T) of *N. meningitidis* (Z-score = 20.1, RMSD = 2.6 Å) (Table 4). NspA is an important adhesin in *N. meningitidis* and binds to hydrophobic ligands, such as lipids. SPAAN analysis, in our study also predicted WP_004684144.1 as an adhesin (P_ad_ value = 0.74). Foldseek has found its best match with OMBB protein of *Legionella rubrilucens*. Based on sequence similarity, PANNZER has predicted its closest match (PPV value 0.78) with Heat resistant agglutinin 1 (Hra1). Heat-resistant agglutinin 1 (Hra1) is an accessory colonization factor of enteroaggregative *E. coli* (EAEC) and can act as an invasin and adhesin.^67^ This suggests that WP_004684144.1 is an important adhesin protein involved in host-pathogen interactions. eggNOG-mapper also predicted it as an ortholog of a surface antigen. Based on the consensus from all four annotation tools, it is reasonable to hypothesize that WP_004684144.1 is a surface-exposed, immunogenic protein, potentially involved in host immune interactions or adhesion mechanisms. No amino acid sequence variation was identified across 46 strains of *B. melitensis*, indicating that the protein is highly conserved and can serve as a promising vaccine candidate (Figure 3, Table S3).

##### 3.3.3.6 WP_004684330.1

WP_004684330.1 is annotated as YadA-like family protein in NCBI but not annotated in UniProt. Structural modelling tools predicted a trimeric structure with each protomer having four β strands that forms a 12-stranded transmembrane β-barrel (Figure 3). A previous in silico study in *B. abortus* identified trimeric autotransporter adhesins (TAAs) such as BatA to be involved in bacterial virulence and adhesion. These TAAs contain an N-terminal YadA-like domain embedded in the OM and a C-terminal α-helical coiled stalk exposed to extracellular environment. The same study traced the evolution of TAAs in different Brucellaceae species, including *B. melitensis*, and predicted a similar YadA-like domain architecture.^68^ BtaE, a YadA-like autotransporter in *Brucella suis*, binds to hyaluronic acid in host cell.^69^ The predicted structure of WP_004684330.1 aligned well with β-barrel architecture of these TAAs by US-align (RMSD = 6.51 Å). Together, these studies highlight the importance of YadA-like autotransporters in *Brucella* virulence. SPAAN predicted WP_004684330.1 to be an adhesin (P_ad_ value = 0.75), suggesting its possible role in host cell attachment, colonization, and pathogenesis. Structural alignment using DALI placed it closest to *Haemophilus influenzae* adhesin Hia (PDB Id: 2GR8; Z-score = 11.8, RMSD = 0.9 Å; Table 4). Hia has a translocator β-barrel domain and an α-helical passenger domain. The passenger domain crosses the outer membrane through the Hia β-barrel to mediate bacterial adhesion to the host respiratory epithelium.^70^ Functional annotation using Foldseek and PANNZER predicted best match with *E. coli* immunoglobin binding protein EibB. EibB binds to the Fc portion of immunoglobulin G (IgG) in a non-immune manner to evade host immune response by avoiding complement activation and phagocytosis.^71^ According to eggNOG-mapper, the protein sequence of WP_004684330.1 is orthologous to the Hep_Hag domain protein. The Hep_Hag domain is a seven amino acid repeat that is found in haemagglutinins and invasins. The Hep_Hag domain is the immunodominant region that represents the minimum epitope required to stimulate an antibody response in *Burkholderia mallei.*^72^ These findings altogether suggest that WP_004684330.1 is a YadA-like (autotransporter adhesin family) protein, involved in host cell attachment and immunoglobulin binding. Six amino acid residues varied across 46 strains of *B. melitensis*: G49, A51, V118, A119, and D120 in α-helical coiled stalk, and R204 in transmembrane β- barrel (Figure 3, Table S3).

#### 3.3.4 Group D

Group D proteins are novel β-barrel proteins identified in this study. Currently these proteins are only listed in NCBI and/or UniProt databases, with no prior experimental or in silico characterization. The proteins are: WP_002968232.1, WP_004682315.1, WP_004684108.1, WP_004684499.1, WP_005970295.1, and WP_075589504.1 (Table 2).

##### 3.3.4.1 WP_002968232.1

WP_002968232.1 is annotated as an OMBB protein in both NCBI and UniProt. Structural model of WP_002968232.1 from all five tools showed a β-barrel architecture consisting of 18 anti-parallel β-strands (Figure 3). SPAAN predicted it as an adhesin-like protein (P_ad_ = 0.51). WP_002968232.1 showed the best structural match with hemolysin activator protein CdiB of *Acinetobacter baumannii* ACICU (PDB Id: 6WIL; Z-score = 16.4, RMSD = 3.9 Å; Table 4). CdiB transporters are members of Omp85 superfamily that form16 stranded β-barrels. CdiB is a part of contact dependent growth inhibition system in bacteria that uses Type Vb secretion mechanism to export toxin CdiA across the OM.^73^ Although architecture of WP_002968232.1 does not align completely with CdiB transporter, its high DALI Z-score suggests that it may function as an OM transporter . Foldseek identified an additional structural homolog - an uncharacterized protein from *Mesorhizobium amorphae.* Sequence-based annotation using PANNZER and eggNOG-mapper confirmed it as OMBB protein. Taken together, these data suggest that WP_002968232.1 is an outer membrane transporter that may mediate protein secretion or contact-dependent inhibition. Sequence comparison of WP_002968232.1 across 46 *B. melitensis* strains revealed variations at only two positions: ICL region (D48) and transmembrane β- barrel (A483) (Figure 3, Table S3).

##### 3.3.4.2 WP_004682315.1

WP_004682315.1 is annotated in NCBI and UniProt as an OM β-barrel domain-containing protein. Structural model showed a β-barrel architecture consisting of eight anti-parallel β-strands (Figure 3). SPAAN identified WP_004682315.1 as a probable adhesin, with a P_ad_ value of 0.67. DALI analysis showed the best structural alignment with *N. meningitidis* NspA (PDB Id: 1P4T; Z-score = 18.3, RMSD = 2.6 Å; Table 4) which plays an important role in bacterial attachment to the host cells and promotes colonization.^74^ Foldseek identified the best structural match of WP_004682315.1 with an uncharacterized protein from *B. abortus*. Sequence-based analysis using PANNZER indicated its closest homolog to be Heat-Resistant Agglutinin 1 (Hra1), an accessory colonization factor in EAEC known to function as an adhesin. These findings suggest that WP_004684144.1 may serve as a key adhesin involved in host-pathogen interactions. eggNOG-mapper annotated it as an ortholog of a surface antigen. This suggests that WP_004682315.1, structural homolog of NspA, might possess adhesin-like properties. Consensus output for this protein from all four annotation tools is similar to WP_004684144.1 present in Group C indicating functional similarity and potential evolutionary relatedness between the two proteins (Section 3.3.3.5). Sequence variation of WP_004682315.1 across 46 strains of *B. melitensis* revealed two variations at position V136 and D247 in ECL region (Figure 3, Table S3).

##### 3.3.4.3 WP_004684108.1

WP_004684108.1 is annotated as alginate export family protein in NCBI but not annotated in UniProt. Structural model generated by five modelling tools revealed a β-barrel architecture comprising 18 anti- parallel β-strands (Figure 3). SPAAN determined that WP_004684108.1 is likely an adhesin, having P_ad_ value of 0.6. WP_004684108.1 showed best structural alignment with alginate production protein AlgE (PDB Id: 3RBH) of *P. aeruginosa* (Z-score of 30.8, RMSD = 2.6 Å) (Table 4). Foldseek matched it to an uncharacterized protein of Cyclobacteriaceae bacterium. Sequence-based analysis (PANNZER and eggNOG-mapper) also predicted it as an alginate export domain-containing protein that forms a passive pore for diffusion of small hydrophilic molecules across OM. AlgE forms an anion-specific channel that exports the exopolysaccharide alginate, a key factor in lung colonization by pathogens in cystic fibrosis patients.^75^ Structural homology with AlgE suggests that WP_004684108.1 might be an Alginate export porin (AEP) family protein and is involved in secretion of virulence factors in *B. melitensis.* Multiple Sequence Alignment of WP_004684108.1 homologs across 46 *B. melitensis* strains showed four variations in β-barrel domain (E209, T429) and ECL region (S242, S383) (Figure 3, Table S3).

##### 3.3.4.4 WP_004684499.1

WP_004684499.1 is annotated in NCBI as autotransporter outer membrane β-barrel. Structural model generated by five modelling tools showed that it has a 12-stranded transmembrane β-barrel domain at the C-terminal, and an N-terminal passenger domain is extended towards the extracellular environment (Figure 3). SPAAN predicted WP_004684499.1 as an adhesin (P_ad_ value = 0.91). It has been identified as an outer membrane β-barrel domain-containing autotransporter by all four annotation tools (DALI, Foldseek, PANNZER, and eggNOG-mapper) (Table 4). WP_004684499.1 exhibited best structural alignment (Z-score = 31.8, RMSD = 2.3 Å) with serine protease EspP (PDB Id: 3SLT) of *E. coli* O157:H7 (Table 4). EspP is a member of serine protease autotransporters of Enterobacteriaceae (SPATE) which is capable of cleaving pepsin A and human coagulation factor V that could contribute to the mucosal haemorrhage in patients with haemorrhagic colitis.^76^ This suggests that WP_004684499.1 may serve similar functions in *B. melitensis*. It was found that the passenger domain of WP_004684499.1 also contains PATR (<u>P</u>assenger <u>A</u>ssociated <u>T</u>ransport <u>R</u>epeat) sequence ‘GLNKSGAGQLTLSGANTYGGATTIDGGVLLQGE’, which is required for efficient export of the passenger domain during growth and infection.^77^ Sequence comparison across 46 strains of *B. melitensis* showed 11 variations in passenger domain at positions M1, G2, E4, N5, K6, V7, P8, R9, S11, G64, G517 (Figure 3, Table S3).

##### 3.3.4.5 WP_005970295.1

WP_005970295.1 is annotated only in NCBI as autotransporter assembly complex family protein but not annotated in UniProt. Structural model constructed from five modelling tools revealed a β-barrel of 16 β-strands, and three POTRA domains in the N-terminal region (Figure 3). It was identified as a homolog of Translocation and assembly module TamA by both structure- and sequence-based annotation tools. TamA has been implicated in translocation of the passenger domains of a subset of bacterial autotransporters.^78^ The C-terminal β-strand of the barrel forms an unusual inward kink, which weakens the lateral barrel wall and creates a gate for substrate access to the lipid bilayer.^79^ The architecture of this protein aligns with the typical structure of TamA in *E. coli* (PDB Id: 4C00; Z score = 40.3, RMSD = 3.8 Å; Table 4), suggesting that *B. melitensis* possesses a translocation and assembly module (TAM) in addition to the BAM complex for outer membrane biogenesis. TamA forms a complex with periplasmic TamB that consists of a concave, taco-shaped β-sheet with a hydrophobic interior.^80^ Studies in *B. suis* have shown that the TamB homolog, known as MapB, controls translocation of a subset of OMPs to the OM.^81^ Our study also identified a TamB homolog (WP_005970293.1), further supporting the presence of a complete translocation and assembly module (TAM) in *B. melitensis.* Amino acid sequence comparison analysis across 46 strains of *B. melitensis* revealed five variations at positions S75, P158, and P294 in β-barrel domain, and S328 and H332 in POTRA domains, indicating that it is mostly a conserved protein with only a few variations (Figure 3, Table S3).

##### 3.3.4.6 WP_075589504.1

WP_075589504.1 is annotated as OmpP1/FadL family transporter in NCBI. Structural model of WP_075589504.1 constructed using five modelling tools comprises 12 anti-parallel β-strands with a prominent lateral gate between β1 and β12 (Figure 3). WP_075589504.1 was predicted to be a putative adhesin by SPAAN, with a P_ad_ value of 0.77. Structural annotation using DALI aligns it with OMP FadL (PDB Id: 3DWO) of *P. aeruginosa* (Z-score = 26.7, RMSD = 2.4 Å) (Table 4). Foldseek matched this protein with an uncharacterized protein of *Candidatus Brocadia caroliniensis*. Sequence-based annotation studies have suggested that this protein is homologous to OMPP1/FadL/TodX family protein, involved in outer membrane transport of hydrophobic molecules. *E. coli* FadL is a well-characterized family of OMPs that transport long-chain fatty acids (LCFAs). These proteins have a 14 stranded β-barrel and use a unique lateral diffusion mechanism, allowing hydrophobic molecules to exit the barrel laterally into the outer membrane.^82^ Although the structure of WP_075589504.1 lacks two β-strands compared to FadL family proteins but the DALI match showed a high Z score. There are evidences which suggest that some 12 stranded β-barrel proteins are also involved in transport of hydrophobic or amphipathic substrates through the barrel wall into the OM. For instance, COG4313 family proteins, such as Pput2725 from *Pseudomonas putida*, structurally differ from FadL by having a 12-stranded β-barrel but share a common transport mechanism of lateral diffusion to transport fatty acids.^83^ This suggests that despite topological differences, WP_075589504.1 could be involved in fatty acid transport using lateral diffusion mechanism and can be a functional analogue of FadL. Sequence variation analysis revealed only one variation in transmembrane β-barrel at position V184 among 46 *B. melitensis* strains (Figure 3, Table S3).

### 3.4 Characterization of OMBB protein complexes

Homologs of three well-known bacterial OMBB proteins - TolC, LptD and BamA - were identified for *B. melitensis* 16M in our study. Since these are part of complexes which span the OM envelope, we attempted to discover these complexes and further studied their structure and PPIs. These complexes include the RND (Resistance-Nodulation-Division) efflux pump, the lipopolysaccharide (LPS) transport (Lpt) complex, and the β-barrel assembly machinery (BAM) complex. In this study, we have successfully identified and characterized these protein complexes that provide critical insights into their molecular architecture and functional dynamics.

#### 3.4.1 RND efflux pumps

TolC is a key component of efflux systems such as AcrAB−TolC in *Enterobacteriaceae*, MexAB−OprM, MexCD−OprJ, and MexXY−OprM in *P. aeruginosa.*^56,57^ In Gram-negative bacteria, TolC along with other proteins forms a tripartite Resistance-Nodulation-Division (RND) pump composed of an inner membrane protein (IMP), a membrane fusion protein (MFP), and an outer membrane factor (OMF), that enables toxic compounds to be expelled out of the cell envelope.^58^ Posadas et al. identified a TolC homologue of *B. suis* that is involved in resistance to antimicrobial compounds and virulence. They studied the *bepC* gene that encodes for TolC family proteins in *B. suis.*^84^ In 2024, a study was conducted to investigate the genetic determinants of antimicrobial resistance (AMR) in 24 *B. melitensis* strains from India, revealing a 16.67% resistance rate to rifampicin and doxycycline. Whole-genome sequencing identified efflux-related genes (mprF, bepC, bepD, bepE, bepF, bepG), but no classical AMR genes were detected.^52^ In this study, we identified Bep proteins (BepC, BepD, BepE, BepF, and BepG) and predicted their structures that were subsequently computationally assembled to determine the feasibility of formation of RND efflux pumps. Sequence anlaysis using BLASTP showed that BepC (WP_002966799.1) is a homolog of TolC family protein, BepD (WP_004685422.1) and BepF (WP_004681660.1) are homologous to efflux RND transporter periplasmic adaptor subunit AcrA (*E. coli*). BepE (WP_004686764.1) and BepG (WP_004686857.1) were found to be homologous to efflux RND transporter permease subunit (*E. coli*). The structure of *E. coli* AcrAB-TolC efflux system ^58^ was used as a reference to predict two tripartite RND efflux pumps in *B. melitensis*: BepDE-BepC and BepFG-BepC (Figure 4). DALI-based structural homology search for BepC predicted closest match with *E. coli* TolC. BepD and BepF showed closest match with *E. coli* AcrA having highest Z score. Similarly, top hit homologs for BepE and BepG were *E. coli* AcrD and *P. aeruginosa* MexY, respectively. Like other Gram-negative bacteria, *B. melitensis* also possess such RND efflux systems that might be responsible for intrinsic resistance against antibiotics. BepC is an OMF that forms a channel across outer membrane and involved in extrusion of toxins, metabolites, antibiotics, and virulent factors. It is a homotrimer that forms α/β-barrel and interacts with periplasmic BepD or BepF. BepD and BepF are hexameric proteins that forms a bridge between OMF and IMP. Further, inner membrane has BepE or BepG existing as homotrimer forming a triangular funnel like structure for translocation of wide range of molecules. The interactions between BepC-BepD, BepD-BepE, BepC-BepF and BepF-BepG were identified using PDBePISA, revealing hydrogen bonds, salt bridges, and hydrophobic contacts among the interfacing residues. The observed interactions exhibited negative ΔG values (ΔG < 0 indicates that the interface formation is thermodynamically favorable, typically hydrophobic in nature, and may imply strong protein-protein affinity), indicating thermodynamically stable associations (Table 6). These findings identified a complete and stable RND efflux channel spanning the *B. melitensis* cell envelope. Insights into structural composition of these pumps and understanding the mechanisms of RND efflux pump-mediated antibiotic resistance can contribute to the development of new strategies to combat antibiotic-resistance in bacteria.

**Table 6:** Protein–protein interactions and free energy changes (ΔG) of predicted protein complexes.

| Protein complex | Interacting Proteins | | $\Delta G$ value (kcal/mol)<sup>†</sup> |
| --- | --- | --- | --- |
|  | Protein 1 | Protein 2 |  |
| RND efflux system | BepC Chain A | BepD Chain A | -4.1 |
|  |  | BepD Chain B | -3.7 |
|  |  | BepD Chain C | -1.2 |
|  | BepE Chain A | BepD Chain A | -7.7 |
|  |  | BepD Chain B | -6.2 |
|  |  | BepD Chain C | -2.9 |
| Lpt Complex | LptD | LptE | -17.6 |
|  |  | LptA | -4.4 |
|  | LptA | LptC | -11.9 |
|  | LptC | LptF | -11.8 |
|  |  | LptG | -18.2 |
|  | LptF | LptG | -17.9 |
|  |  | LptB | -13.8 |
|  | LptG | LptB | -17.6 |
|  | LptB | LptB | -2.6 |
| **Bam Complex** | BamA | BamD | -5 |
|  |  | BamE | -12.1 |
|  | BamD | BamE | -7.9 |

$\dagger\Delta G$ indicates the solvation free energy gain upon formation of the interface, in kcal/mol, predicted by PDBePISA. Negative $\Delta G$ corresponds to hydrophobic interfaces, or positive protein affinity.

#### 3.4.2 Lpt Complex

In *E. coli*, seven protein components form the Lpt machinery: LptA, LptB, LptC, LptD, LptE, LptF, and LptG. The Lpt machinery is composed of two sub-assemblies. In the outer membrane, LptD forms β-barrel structure and associates with the lipoprotein, which functions as a plug within the complex. LptD plays a key role in the final step of LPS insertion into the outer membrane, ensuring proper membrane biogenesis and integrity. At the inner membrane, an ABC transporter is present, consisting of the cytoplasmic dimeric LptB along with the transmembrane polytopic proteins LptF and LptG. LptB₂FG is an ABC transporter in which LptB act as molecular motor for ATP hydrolysis and both LptF and LptG are polytopic transmembrane proteins, meaning they span the membrane multiple times. They contain six transmembrane helices each, forming a channel-like structure within the inner membrane. This LptB_2_FG complex interacts with LptC, an auxiliary inner membrane protein. Connecting these two subassemblies is the periplasmic lipoprotein LptA that adopts a β-jellyroll fold, forming an elongated structure that facilitates its role as a molecular bridge. LptA contains a hydrophobic groove that interacts with the acyl chains of LPS, allowing for efficient transport.^85^ In this study we identified the components of Lpt complex and assembled their 3D models spanning the bacterial membrane. These annotated components include: LptA (WP_004686649.1), LptB (WP_002971717.1), LptC (WP_004684681.1), LptD (WP_004683433.1), LptE (WP_002964883.1), LptF (WP_004683435.1), and LptG (WP_002967526.1). AlphaFold 3 generated models of LptDE and LptB_2_FGC were predicted to have a good ranking score of 0.86 and 0.79, respectively. To validate the formation of a functional complex, interactions between β jelly roll of LptD-LptA and LptA-LptC were analyzed, that forms a continuous channel across the periplasm for transport of LPS molecules to the outer membrane (Figure 5). Interactions involve hydrogen bonds, salt bridges and hydrophobic contacts between the interfacing residues with negative ΔG value indicating formation of a thermodynamically stable complex (Table 6).

#### 3.4.3 BAM Complex

Bam complex of *E. coli* comprises five components: BamA, BamB, BamC, BamD, and BamE. In contrast to this, *B. melitensis* encodes homologs of only BamA, BamD, and BamE proteins, while BamB and BamC proteins are absent.^18,65^ In this study, we have identified the constituent proteins of the BAM complex in *B. melitensis* and analyzed their interactions fundamental to the assembly and maintenance of the bacterial outer membrane. Bam complex of *B. melitensis* consists of a key component BamA (WP_004683866.1) that interacts with two periplasmic lipoproteins, BamD (WP_002964530.1) and BamE (WP_005969315.1). Here, we have assembled the BAM complex in *B. melitensis* membrane (Figure 6A) and compared it with *E. coli* BAM complex. Construction of Bam complex using AlphaFold 3 gave a high confidence model with iptm score of 0.77 and ranking score of 0.83.

The structure of the fully assembled BAM complex of *E. coli* has been resolved using X-ray crystallography at 3.4 Å resolution by Bakelar et al. in 2015. They have identified essential amino acid residues in BamA that interact with BamD and BamE.^86^ In 2021, Hawley et al. predicted structural model of protein complexes in *Treponema pallidum* and studied their structural characteristics by comparing them with homologous proteins of other well-known proteobacteria using structure-based sequence alignment.^87^ Similarly, structure based sequence alignment of *B. melitensis* BamA with *E. coli* BamA was performed to reveal the equivalent amino acid residues in *B. melitensis* BamA that might interact with BamD and BamE using PROMALS3D. PROMSLS3D constructs alignments for multiple protein structures using information from available 3D structures, database homologs, and predicted secondary structures. Table S4 shows equivalent residues that have been shown on the structural model of *B. melitensis* BamA (Figure 6B, 6C, 6E, 6F). BamD majorly interacts with POTRA 5 and POTRA 2 of BamA. BamE interacts with POTRA 5 and periplasmic loop 3 of BamA towards periplasmic side. PDBePISA predicted all the residues in BamA present at the interface of BamADE (Figure 6D, 6G) complex, and the results aligned with results of PROMALS3D. These equivalent residues are present at interface and involved in formation of hydrogen bonds, salt bridges, and hydrophobic interactions, resulting in thermodynamically stable complex with negative ΔG values (Table 6). Interaction of *B. melitensis* BamA with BamDE involves more contact residues as compared to *E. coli* BamA, which may be due to lack of BAM accessory subunits (BamB and BamC) in *B. melitensis.* This strongly suggests that *B. melitensis* may utilize a distinct mechanism for transferring nascent OMPs to the β-barrel of BamA, with these interactions potentially compensating for the absence of BamB and BamC.

## 4. Conclusion

*B. melitensis*, like many Gram-negative bacteria, relies on its outer membrane to survive and adapt within the host environment. This study provides a robust computational framework for *B. melitensis* OMBB identification and validation, followed by comprehensive in silico characterization of OMBB proteins and their complexes. Our results offer important insights into the structural architecture, potential functions, and roles in membrane assembly and transport. By integrating structural models, membrane insertion analyses, and PPI modelling, we successfully identified key OMBB proteins, their associated complexes, and their possible amino acid interactions. This study elucidates the architecture of *B. melitensis* outer membrane by presenting a molecular view of protein complexes. PPIs within multi-protein complexes highlight the sophisticated outer membrane machinery involved in OM biogenesis and efflux of toxic substances. Most of the OMBB proteins were found to be highly conserved and can also serve as promising targets for diagnostics, vaccine development, and therapeutic designs.

## Acknowledgements

JK is a recipient of junior research fellowship from the University Grants Commission, Government of India. AP is a recipient of junior research fellowship from the Department of Biotechnology, Government of India.

## Author contributions statement

**Jahnvi Kapoor:** methodology, software, investigation, data analysis, data curation, writing— original draft preparation. **Amisha Panda:** data analysis, visualization, writing—reviewing and editing. **Ilmas Naqvi**: visualization, writing—reviewing and editing. **Satish Ganta:** visualization, writing—reviewing and editing. **Sanjiv Kumar:** conceptualization, methodology, software, writing—reviewing and editing, supervision. **Anannya Bandyopadhyay**: methodology, data analysis, writing—original draft preparation, writing—reviewing and editing, supervision. All authors reviewed the final draft of the manuscript. All authors contributed to the manuscript revision, read and approved the submitted version.

## Data Availability Statement

The data supporting this study’s findings are available from the corressponding author upon reasonable request.

