## Supplementary material for "Identification and Characterization of Outer Membrane Proteins and Membrane Spanning Protein Complexes in *Brucella melitensis*"

**Table S1:** Comprehensive results from computational tools employed in our study to predict outer membrane β-barrel proteins

| **Protein Accession No.** | **Protein name**^†^ | **OMPdb** | **TMBETADISC** | **MCMBB** | **TMbed** | | **Pepstats** | | **PSORTb** | | **SignalP** | | **SPAAN** | **CELLO** |
| --- | --- | --- | --- | --- | --- | --- | --- | --- | --- | --- | --- | --- | --- | --- |
|  |  |  |  |  | **tmbed_b** | **tmbed_B** | **AA Length** | **MolWt (Da)** | **Localization** | **Score** | **Signal Peptide** | **CS Position** | **P_ad_ Value** |  |
| **Group A** |  |  |  |  |  |  |  |  |  |  |  |  |  |  |
| WP_002964666.1 | OmpW family protein | 500 | Outer Membrane Protein | 0.035 | 39 | 37 | 227 | 24500.98 | Outer Membrane | 10 | SP(Sec/SPI) | CS pos: 23-24. AFA-AD. Pr: 0.9846 | 0.682036 | Periplasmic |
| WP_002965367.1 | 25 kDa outer membrane protein | 500 | Outer Membrane Protein | 0.07 | 35 | 33 | 230 | 24748.68 | Outer Membrane | 10 | SP(Sec/SPI) | CS pos: 22-23. AYA-AD. Pr: 0.9914 | 0.799603 | Outer  Membrane |
| WP_002965368.1 | 25 kDa outer membrane protein | 500 | Outer Membrane Protein | 0.062 | 35 | 33 | 228 | 24586.35 | Outer Membrane | 10 | SP(Sec/SPI) | CS pos: 22-23. ANA-AD. Pr: 0.9819 | 0.597688 | Outer  Membrane |
| WP_004683466.1 | outer membrane protein Omp25 | 500 | Outer Membrane Protein | 0.063 | 35 | 33 | 213 | 23185.12 | Outer Membrane | 10 | SP(Sec/SPI) | CS pos: 23-24. AFA-AD. Pr: 0.9862 | 0.574148 | Outer  Membrane |
| WP_004683739.1 | 25 kDa outer membrane protein | 500 | Outer Membrane Protein | 0.055 | 35 | 33 | 236 | 25242.25 | Outer Membrane | 10 | SP(Sec/SPI) | CS pos: 22-23. AKA-AD. Pr: 0.9781 | 0.663293 | Outer  Membrane |
| WP_004681766.1 | outer membrane protein Omp31 | 501 | Outer Membrane Protein | 0.06 | 36 | 35 | 240 | 25323.2 | Outer Membrane | 10 | SP(Sec/SPI) | CS pos: 19-20. AMA-AD. Pr: 0.9911 | 0.835538 | Outer  Membrane |
| WP_002970988.1 | porin | 500 | Outer Membrane Protein | 0.042 | 73 | 62 | 367 | 39478.34 | Outer Membrane | 9.93 | SP(Sec/SPI) | CS pos: 22-23. AQA-AD. Pr: 0.9805 | 0.594483 | Outer Membrane |
| WP_006256196.1 | porin | 500 | Outer Membrane Protein | 0.039 | 71 | 62 | 362 | 38704.63 | Outer Membrane | 9.93 | SP(Sec/SPI) | CS pos: 22-23. AQA-AD. Pr: 0.9808 | 0.602086 | Outer  Membrane |
| **Group B** |  |  |  |  |  |  |  |  |  |  |  |  |  |  |
| WP_002966799.1 | multidrug efflux RND transporter outer membrane | 500 | Outer Membrane Protein | 0.046 | 18 | 18 | 456 | 48490.71 | Outer Membrane | 10 | SP(Sec/SPI) | CS pos: 28-29. GQA-AL. Pr: 0.3201 | 0.443155 | Outer  Membrane |
| WP_004681095.1 | TonB-dependent receptor domain-containing protein | 500 | Outer Membrane Protein | 0.039 | 96 | 88 | 661 | 72932.84 | Outer Membrane | 9.93 | SP(Sec/SPI) | CS pos: 23-24. SQA-QE. Pr: 0.9745 | 0.712876 | Outer Membrane |
| WP_004683979.1 | TonB-dependent receptor | 500 | Outer Membrane Protein | 0.041 | 102 | 93 | 620 | 66853.9 | Outer Membrane | 10 | SP(Sec/SPI) | CS pos: 23-24. ALA-QD. Pr: 0.9904 | 0.696891 | Outer Membrane |
| WP_005972352.1 | TonB-dependent copper receptor | 500 | Outer Membrane Protein | 0.037 | 103 | 92 | 676 | 75161.78 | Outer Membrane | 10 | SP(Sec/SPI) | CS pos: 32-33. AYA-QE. Pr: 0.9937 | 0.432338 | Outer Membrane |
| **Group C** |  |  |  |  |  |  |  |  |  |  |  |  |  |  |
| WP_002971090.1 | hypothetical protein | 42 | Outer Membrane Protein | 0.021 | 48 | 47 | 267 | 30043.87 | Unknown | 2.5 | SP(Sec/SPI) | CS pos: 20-21. APA-AD. Pr: 0.8858 | 0.730601 | Outer Membrane |
| WP_004683433.1 | LPS-assembly protein LptD | 500 | Outer Membrane Protein | 0.026 | 118 | 105 | 792 | 88474.43 | Outer Membrane | 10 | SP(Sec/SPI) | CS pos: 36-37. AQA-QD. Pr: 0.9154 | 0.644231 | Outer Membrane |
| WP_004683866.1 | outer membrane protein assembly factor BamA | 500 | Outer Membrane Protein | 0.04 | 73 | 70 | 781 | 85918.96 | Outer Membrane | 10 | SP(Sec/SPI) | CS pos: 38-39. AEA-AV. Pr: 0.3549 | 0.411397 | Outer Membrane |
| WP_004683944.1 | outer membrane protein | 500 | Outer Membrane Protein | 0.052 | 37 | 34 | 212 | 22108.63 | Outer Membrane | 9.93 | SP(Sec/SPI) | CS pos: 24-25. AFA-AD. Pr: 0.9962 | 0.823197 | Extracellular |
| WP_004684144.1 | outer membrane protein | 500 | Outer Membrane Protein | 0.041 | 40 | 35 | 274 | 29184.35 | Unknown | 2.5 | SP(Sec/SPI) | CS pos: 26-27. ASA-TD. Pr: 0.8883 | 0.742591 | Outer Membrane |
| WP_004684330.1 | YadA-like family protein | 510 | Outer Membrane Protein | 0.052 | 16 | 15 | 227 | 23599.61 | Cytoplasmic Membrane | 9.86 | SP(Sec/SPI) | CS pos: 27-28. AKA-EE. Pr: 0.8177 | 0.75465 | Extracellular |
| **Group D** |  |  |  |  |  |  |  |  |  |  |  |  |  |  |
| WP_002968232.1 | outer membrane beta-barrel protein | 500 | Outer Membrane Protein | 0.032 | 87 | 81 | 512 | 55085.26 | Outer Membrane | 9.52 | SP(Sec/SPI) | CS pos: 36-37. AYA-QD. Pr: 0.9916 | 0.515837 | Outer Membrane |
| WP_004682315.1 | outer membrane protein | 500 | Outer Membrane Protein | 0.009 | 40 | 37 | 284 | 31493.17 | Outer Membrane | 10 | SP(Sec/SPI) | CS pos: 23-24. ASA-AD. Pr: 0.9437 | 0.6742 | Outer Membrane |
| WP_004684108.1 | alginate export family protein | 252 | Outer Membrane Protein | 0.034 | 84 | 81 | 586 | 63993.59 | Outer Membrane | 9.52 | SP(Sec/SPI) | CS pos: 30-31. AMA-QD. Pr: 0.9030 | 0.600102 | Outer Membrane |
| WP_004684499.1 | autotransporter outer membrane beta-barrel | 268 | Outer Membrane Protein | 0.064 | 62 | 59 | 1440 | 146738.6 | Outer Membrane | 9.82 | OTHER |  | 0.91096 | Extracellular |
| WP_005970295.1 | autotransporter assembly complex family protein | 500 | Outer Membrane Protein | 0.017 | 74 | 69 | 639 | 68866.99 | Outer Membrane | 10 | SP(Sec/SPI) | CS pos: 18-19. ALA-FE. Pr: 0.8634 | 0.394226 | Outer Membrane |
| WP_075589504.1 | OmpP1/FadL family transporter | 500 | Outer Membrane Protein | 0.035 | 58 | 51 | 323 | 35148.34 | Outer Membrane | 9.49 | OTHER |  | 0.770266 | Outer Membrane |

†Protein names from NCBI, searched using Protein Accession no.

**Table S2:** 46 strains of *B. melitensis* searched for predicted OMP’s variations compared to the reference genome 16M

| **Assembly Accession** | **Strain Name** | **Genome Size (Mb)** | **CDS**^†^ |
| --- | --- | --- | --- |
| GCF_000007125.1 | 16M | 3.295 | 2943 |
| GCF_000022625.1 | ATCC 23457 | 3.311 | 3010 |
| GCF_000192725.1 | M28 | 3.312 | 3003 |
| GCF_000192885.1 | M5-90 | 3.312 | 2990 |
| GCF_000227645.1 | NI | 3.294 | 2972 |
| GCF_000740355.1 | ether | 3.311 | 2988 |
| GCF_001307475.2 | C-573 | 3.311 | 2961 |
| GCF_001431745.1 | 20236 | 3.312 | 3009 |
| GCF_001715485.1 | 2.01E+09 | 3.31 | 2987 |
| GCF_002191235.1 | BwIM_AFG_63 | 3.311 | 3010 |
| GCF_002191295.1 | BwIM_IRN_37 | 3.313 | 3016 |
| GCF_002191335.1 | BwIM_IRQ_32 | 3.312 | 3014 |
| GCF_002191355.1 | BwIM_ITA_45 | 3.31 | 2991 |
| GCF_002191375.1 | BwIM_ITA_55 | 3.31 | 2990 |
| GCF_002191455.1 | BwIM_SYR_04 | 3.313 | 3014 |
| GCF_002191575.1 | BwIM_TKM_56 | 3.311 | 3012 |
| GCF_002191615.1 | BwIM_TUR_03 | 3.313 | 3012 |
| GCF_002191655.1 | BwIM_TUR_17 | 3.313 | 3012 |
| GCF_002191675.1 | BwIM_TUR_19 | 3.313 | 3016 |
| GCF_002191755.1 | BwIM_TUR_59 | 3.313 | 3015 |
| GCF_002191915.1 | BwIM_SAU_09 | 3.311 | 3008 |
| GCF_002192095.1 | BwIM_SYR_26 | 3.313 | 3015 |
| GCF_002192155.1 | BwIM_TUR_39 | 3.313 | 3015 |
| GCF_002214285.1 | QY1 | 3.311 | 2897 |
| GCF_002262955.1 | BY38 | 3.312 | 3011 |
| GCF_002263015.1 | BL | 3.313 | 3014 |
| GCF_002763615.1 | B.melitensis QH61 | 3.312 | 3009 |
| GCF_002895105.1 | CIIMS-BH-2 | 3.311 | 3007 |
| GCF_002895125.1 | CIIMS-PH-3 | 3.311 | 2994 |
| GCF_002953595.1 | Rev.1 (passage 101) | 3.299 | 2990 |
| GCF_003205535.1 | CIIMS-NV-1 | 3.312 | 3006 |
| GCF_003516045.1 | CIT21 | 3.311 | 3010 |
| GCF_003516065.1 | CIT31 | 3.312 | 3016 |
| GCF_003516085.1 | CIT43 | 3.312 | 3011 |
| GCF_003856415.1 | BmWS93 | 3.312 | 3008 |
| GCF_004208655.1 | B29 | 3.312 | 3011 |
| GCF_004208675.1 | B15 | 3.312 | 3015 |
| GCF_004208695.1 | B9 | 3.312 | 3011 |
| GCF_008761595.1 | M1981 | 3.312 | 3015 |
| GCF_008761615.1 | RM57 | 3.306 | 3001 |
| GCF_009017395.1 | VB12455 | 3.312 | 2854 |
| GCF_023796775.1 | 6144 | 3.312 | 3007 |
| GCF_024802385.1 | PB150210 | 3.311 | 3013 |
| GCF_027625455.1 | TZ | 3.312 | 3014 |
| GCF_036320855.1 | B-HB-9 | 3.306 | 2991 |
| GCF_036967195.1 | ARQ-070 | 3.298 | 2996 |
| GCF_038420175.1 | Rev.1 | 3.299 | 2981 |

†CDS: Coding Sequences

**Table S3:** Sequence variations among 46 ­strains of *B. melitensis*

| **Protein Accession No.** | **Protein name**^†^ | **Total no. of variations** | **Amino acid residues** |
| --- | --- | --- | --- |
| **Group A** |  |  |  |
| WP_002964666.1 | OmpW family protein | 2 | A18, T224 |
| WP_002965367.1 | outer membrane protein | 2 | N91, G175 |
| WP_002965368.1 | outer membrane protein | 0 | - |
| WP_004683466.1 | outer membrane protein Omp25 | 1 | A139 |
| WP_004683739.1 | outer membrane protein | 3 | N2, V13, R57 |
| WP_004681766.1 | outer membrane protein Omp31 | 2 | A7, T176 |
| WP_002970988.1 | porin | 56 | P31, M91, F92, N93, N95, G104, Y106, Q108, T114, S115, R119, H120, Q123, D126, F127, S128, D129, D130, R131, D132, V133, A134, D135, G136, G137, V138, T140, G141, D143, L144, Q145, T150, F154, K155, V179, A180, E210, D211, V212, D213, N214, D215, T217, H224, T253, R264, N280, A296, F298, I299, P301, E302, T305, Q327, I334, D349 |
| WP_006256196.1 | porin | 53 | P31, I55, G85, R91, V92, S93, G95, K104, F106, E108, A114, A115, G119, V120, K123, N126, E127, T128, S130, V133, M134, E135, Q140, L144, R145, I169, S170, S180, D200, N201, D202, G203, G204, Y205, T206, G207, T208, T209, N210, H212, D219, A248, Q259, D275, L291, Y293, Q294, T296, Q297, A300, E322, V329, N344 |
| **Group B** |  |  |  |
| WP_002966799.1 | multidrug efflux RND transporter outer membrane | 0 | - |
| WP_004681095.1 | TonB-dependent receptor domain-containing protein | 0 | - |
| WP_004683979.1 | TonB-dependent receptor | 1 | W315 |
| WP_005972352.1 | TonB-dependent copper receptor | 1 | N538 |
| **Group C** |  |  |  |
| WP_002971090.1 | hypothetical protein | 0 | - |
| WP_004683433.1 | LPS-assembly protein LptD | 5 | Q45, E215, Q292, G347, R554 |
| WP_004683866.1 | outer membrane protein assembly factor BamA | 1 | V318 |
| WP_004683944.1 | outer membrane protein | 1 | N36 |
| WP_004684144.1 | outer membrane protein | 0 | - |
| WP_004684330.1 | YadA-like family protein | 6 | G49, A51, V118, A119, D120, R204 |
| **Group D** |  |  |  |
| WP_002968232.1 | outer membrane beta-barrel protein | 2 | D48, A483 |
| WP_004682315.1 | outer membrane protein | 2 | V136, D247 |
| WP_004684108.1 | alginate export family protein | 4 | E209, S242, S383, T429 |
| WP_004684499.1 | autotransporter outer membrane beta-barrel | 11 | M1, G2, E4, N5, K6, V7, P8, R9, S11, G64, G517 |
| WP_005970295.1 | autotransporter assembly complex family protein | 5 | S75, P158, P294, S328, H332 |
| WP_075589504.1 | OmpP1/FadL family transporter | 1 | V184 |

†Protein names from NCBI, searched using Protein Accession no.

**Table S4:** Equivalent residues in *B. melitensis* BamA identified through structure-based sequence alignment and types of interactions within the Bam complex

| Residues in *E. coli* BamA interacting with BamDE | Equivalent interfacing residues in *B. melitensis* BamA  **(PROMALS3D, PDBePISA)** | Type of interactions  **(PDBePISA)** |
| --- | --- | --- |
| A95 | N110 | Hydrogen Bond |
| S96 | N111 | Interfacing Residue |
| R120 | K132 | - |
| V121 | P133 | Interfacing Residue |
| G122 | R134 | Hydrogen Bond |
| E123 | A135 | Interfacing Residue |
| Y348 | Y361 | Interfacing Residue, Hydrogen Bond |
| I352 | I365 | Hydrogen Bond |
| N357 | N370 | Hydrogen Bond |
| D358 | D371 | Interfacing Residue |
| T359 | K372 | Hydrogen Bond, Interfacing Residue |
| D362 | D375 | Hydrogen Bond |
| R366 | R379 | Hydrogen Bond |
| E373 | E386 | Hydrogen bond, Salt bridge |
| G374 | G387 | Interfacing Residue |
| W376 | A389 | Hydrogen Bond |
| S408 | - | - |
| D410 | D422 | Hydrogen Bond |
| Q411 | Q423 | Interfacing Residue |
| D481 | Y497 | Interfacing Residue |
| P518 | P530 | Hydrogen Bond |
| E521 | D533 | Hydrogen Bond |

**Interfacing residues** are involved in hydrophobic or Van der Waals interactions.

**Table S5:** Amino acid residues involved in protein-protein interactions in predicted protein complexes identified using PDBePISA

| **Protein Complexes** | **Interacting proteins** | **Amino acid Residues involved in hydrogen bonding** | **Interfacing Residues (hydrophobic interactions)** | **Amino acid residues forming salt bridges** |
| --- | --- | --- | --- | --- |
| **RND efflux system** | **BepC-BepD** |  |  |  |
|  | BepC chain A | G176, R167, V175, G380, V379 | V175, E177, G178, T179, R180, T181, E164, A168, A171, R172, G176, E177, V379, Q381, R382, T383, T384, L385, D371, G372, E375, E376, G380, Q381 |  |
|  | BepD chain A | A139, A140 | K135, V136, Q137, S138 |  |
|  | BepD chain B | R127, Q134, A139, A140 | D124, A128, Q130, L131, K135, V136, Q137, S138 |  |
|  | BepD chain C | R127 | A128, Q130, L131, Q134, V136, Q137 |  |
| **RND efflux system** | **BepE-BepD** |  |  |  |
|  | BepE chain A | R190, D192, A653, I655, Y723, E774, D778, S791, S793, D798, R805, R253, S256, N258 | R193, A195, Q196, N198, R526, G527, F530, R534, R555, V574, L575, P576, P577, R583, N652, K656, D657, G658, M659, E690, Y719, P721, K731, M735, R777, P779, K783, H784, F786, A789, M794, I795, P796, A799, I806, V807, L911, D914, L915, D728, E730, K731, A734, M735, W892, K1028, T146, D147, R148, Y149, D150, R193, Q196, Y197, N198, S254, S255, Q257, A259, A260, T261, L262, K265, D266, P320, W364, K500, H227, F231, Y233, S256 | D192, E774, D778 |
|  | BepD chain A | T219, G250, E271, R295, K311, Q316, E342, K366 | L3, N4, I7, V18, F19, A21, Q38, V39, F40, T52, Y53, E54, Y55, A56, R58, S217, D220, E222, D251, T270, T272, G273, T274, G291, Q292, A312, L314, M315, S317, A318, Q321, R341, L343, K344, W347, T361, E362, G363, V364, I365, V376 | R295, K311 |
|  | BepD chain B | R58, S217, E271 | M1, N4, A56, A57, N215, F216, T219, D220, T221, E222, L268, D269, T270, T272, G273, T274, G276, F293, M315, Q316, S317, A318, F322, Y324, V334, V364, I365, V368, P369 |  |
|  | BepD chain C |  | L3, I7, A56, R58, F293, R295 |  |
| **Lpt Complex** | **LptD-LptA** |  |  |  |
|  | LptD | L16, C18, L20, A21, P23, V25, V27, L30, Q37, A56, L59 | A11, R12, G13, T14, A15, A17, V19, L22, F24, S26, A28, S32, P33, A36, A39, A42, L54, D57, D86 | A21, V27 |
|  | LptA | A100, A128, V131, G145, Y147, A149, N151, Q159, K160, V162, T164, D167 | S98, K99, K101, D126, K130, Y132, L133, D143, T144, R146, D148, K150, V161, L163, D165, G166, N168, I169 | A128, Q159 |
| **Lpt Complex** | **LptA-LptC** |  |  |  |
|  | LptA | G20, S22, V23, L24, A25, L26, A27, F29, A31, P32 | R15, T16, L17, R18, I19, F21, A28, A30, Q38, P46, A73, V74, F75, T76, G77, N78, V79, A80, V81, G84, D85, G115 | V23, L26, P32 |
|  | LptC | E137, K214, L216, F218, E219, Y220, V222, H223, M224, V226 | G136, N169, G170, L171, I195, N197, G198, T199, T200, I202, G213, V215, D227, G228, N229, T230, L231, S232, N234, K235 | E137, E219, H223 |
| **Bam Complex** | **BamA-BamD** |  |  |  |
|  | BamA | R51, D53, Q115, K118, R174, N176, R364, I367, G369 | V52, T56, D59, N60, R64, A83, M84, L86, E105, R106, V108, V109, L113, F114, D122, P136, R166, V168, L170, G171, Q172, V178, I362, Q363, E366, R368, T373, R374, Y376, R380, N385, D388, Q423, L465, G496 | R364 |
|  | BamD | N32, I35, Y40, E42, Q76, Y79, T80, R109, T139, R140, D141 | S30, K31, D34, L37, V41, I44, A71, D74, R75, H77, P78, E81, L87, A90, E102, M106, T112, L113, Y114, P115, T116, P136, D137, A143, R146, Q185, I186, Y189, R193, E195, L197, A198, K201, G205, E209 | E183, R202 |
| **Bam Complex** | **BamA-BamE** |  |  |  |
|  | BamA | K121, A326, R359, R374, N385, D388, R397, L465, L495, R498, I531 | W219, L220, R222, R299, L300, K302, R304, E323, G327, S328, G329, Y330, A331, F332, Q356, D371, D383, G387, M394, R401, E420, P421, S434, G436, F438, I440, I459, E461, F464, R467, I471, I473, G496, L499, L529, T532, N534, F535, Y583, L596, F751, F781 | R359, R374, R397 |
|  | BamE | E40, T43, E44, G45, Y46, V47, T71, Q95, F96 | C30, T31, T32, S34, T35, L36, N37, P38, T41, L42, A52, A68, L69, G70, P72, S73, K89, R90, R92, A94 | E40, D49 |

**Table S6:** Proteins predicted as outer membrane β-barrels by three out of four tools in *B. melitensis* 16M

| **Protein Accession No.** | **NCBI Protein name** | **UniProt Protein name** | **OMPdb Match** | **MCMBB Score** | **TMBETADISC** | **TMbed** |
| --- | --- | --- | --- | --- | --- | --- |
| WP_002963577.1 | enoyl-ACP reductase FabI | Enoyl-[acyl-carrier-protein] reductase [NADH] | + | + | + | - |
| WP_002963579.1 | cold-shock protein | Cold-shock protein | + | + | + | - |
| WP_002963616.1 | acyl carrier protein [Hyphomicrobiales]. | Acyl carrier protein AcpP | + | + | + | - |
| WP_002964019.1 | cell division endopeptidase DipM | - | + | + | + | - |
| WP_002964124.1 | hypothetical protein | Uncharacterized protein | - | + | + | + |
| WP_002964621.1 | cold-shock protein | Cold-shock protein | + | + | + | - |
| WP_002964760.1 | tonB-system energizer ExbB | Biopolymer transport protein ExbB | + | + | + | - |
| WP_002964765.1 | efflux RND transporter periplasmic adaptor subunit | Efflux transporter, RND family, MFP subunit | + | + | + | - |
| WP_002964792.1 | Holliday junction branch migration protein RuvA | Holliday junction branch migration complex subunit RuvA | + | + | + | - |
| WP_002964802.1 | peptidoglycan -binding protein | OmpA/MotB domain protein | + | + | + | - |
| WP_002965036.1 | electron transfer flavoprotein subunit alpha/FixB | Electron transfer flavoprotein subunit alpha | + | + | + | - |
| WP_002965233.1 | enoyl-ACP reductase FabI | Enoyl-[acyl-carrier-protein] reductase [NADH] | + | + | + | - |
| WP_002965683.1 | tetratricopeptide repeat protein | GlcNAc transferase | + | + | + | - |
| WP_002966947.1 | peptidoglycan-associated lipoprotein Pal | Peptidoglycan-associated lipoprotein | + | + | + | - |
| WP_002967320.1 | polysaccharide biosynthesis/export family protein | Polysaccharide export protein | + | + | + | - |
| WP_002970964.1 | Ig-like domain-containing protein | Peptidoglycan-binding LysM | + | + | + | - |
| WP_002970984.1 | phage tail tape measure protein | Phage tail protein | + | + | + | - |
| WP_002971258.1 | HlyD family secretion protein | HlyD family secretion protein | + | + | + | - |
| WP_004681011.1 | autotransporter-associated beta strand | - | + | + | + | - |
| WP_004681262.1 | molybdenum ABC transporter ATP-binding protein | Molybdenum import ATP-binding protein ModC | + | + | + | - |
| WP_004681310.1 | HlyD family secretion protein | Multidrug resistance protein a | + | + | + | - |
| WP_004681382.1 | flagellar basal body L-ring protein FlgH | Flagellar L-ring protein | - | + | + | + |
| WP_004681660.1 | multidrug efflux RND transporter periplasmic adaptor | - | + | + | + | - |
| WP_004681746.1 | exopolysaccharide transport family protein | Succinoglycan biosynthesis transport protein exop | + | + | + | - |
| WP_004681856.1 | GntR family transcriptional regulator | Transcriptional regulator, GntR family protein | + | + | + | - |
| WP_004681901.1 | Asp-tRNA(Asn)/Glu-tRNA(Gln) amidotransferase subunit | Glutamyl-tRNA(Gln) amidotransferase subunit A | + | + | + | - |
| WP_004682166.1 | efflux RND transporter permease subunit | Acriflavin resistance protein f | + | + | + | - |
| WP_004682168.1 | efflux RND transporter periplasmic adaptor subunit | Efflux transporter, RND family, MFP subunit | + | + | + | - |
| WP_004682822.1 | efflux RND transporter periplasmic adaptor subunit | Efflux transporter, RND family, MFP subunit | + | + | + | - |
| WP_004683321.1 | DegQ family serine endoprotease | Probable periplasmic serine endoprotease DegP-like | + | + | + | - |
| WP_004683696.1 | SPOR domain-containing protein | - | + | + | + | - |
| WP_004683827.1 | single-stranded DNA-binding protein | Single-stranded DNA-binding protein | + | + | + | - |
| WP_004684003.1 | DegQ family serine endoprotease | Probable periplasmic serine endoprotease DegP-like | + | + | + | - |
| WP_004684148.1 | tol-pal system protein YbgF | - | + | + | + | - |
| WP_004684153.1 | tetratricopeptide repeat protein | Sel1 repeat family protein | + | + | + | - |
| WP_004684526.1 | ABC transporter substrate-binding protein | - | + | + | + | - |
| WP_004685422.1 | multidrug efflux RND transporter periplasmic adaptor | - | + | + | + | - |
| WP_004685697.1 | DegQ family serine endoprotease | Probable periplasmic serine endoprotease DegP-like | + | + | + | - |
| WP_004685732.1 | HlyD family secretion protein | HlyD family secretion protein | + | + | + | - |
| WP_004686432.1 | recombinase family protein | Resolvase | + | + | + | - |
| WP_004686831.1 | GumC family protein | Succinoglycan biosynthesis transport protein exop | + | + | + | - |
| WP_004686888.1 | HlyD family secretion protein | Fusaric acid resistance protein fuse | + | + | + | - |
| WP_004686962.1 | OmpA family protein | Cell envelope biogenesis protein OmpA | + | + | + | - |
| WP_005970024.1 | recombinase family protein | - | + | + | + | - |
| WP_005972432.1 | efflux RND transporter periplasmic adaptor subunit | Acriflavin resistance protein a | + | + | + | - |
| WP_020699915.1 | LysR family transcriptional regulator | LysR family transcriptional regulator | + | + | + | - |
| WP_041594647.1 | efflux RND transporter periplasmic adaptor subunit | - | + | + | + | - |
| WP_244391821.1 | autotransporter-associated beta strand | - | + | + | - | + |

+ represents positive output given by the computational tool utilised,

- represents negative result given by the computational tool utilised.
